# Redundant cytokine requirement for intestinal microbiota-induced Th17 cell differentiation in draining lymph nodes

**DOI:** 10.1101/2020.04.21.053934

**Authors:** Teruyuki Sano, Takahiro Kageyama, Victoria Fang, Ranit Kedmi, Jhimmy Talbot, Alessandra Chen, Reina Kurakake, Yi Yang, Charles Ng, Susan R. Schwab, Dan R. Littman

## Abstract

Differentiation of intestinal T helper 17 (Th17) cells, which contribute to mucosal barrier protection from invasive pathogens, is dependent on colonization with distinct commensal bacteria. Segmented filamentous bacteria (SFB) are sufficient to support Th17 cell differentiation in mouse, but the molecular and cellular requirements for this process remain incompletely characterized. Here we show that intestine-draining mesenteric lymph nodes (MLN) are the dominant site of SFB-induced intestinal Th17 cell differentiation. Subsequent migration of these cells to the intestinal lamina propria is dependent on their up-regulation of integrin β7. Stat3-dependent induction of RORγt, the Th17 cell-specifying transcription factor, largely depends on IL-6, but signaling through the receptors for IL-21 and IL-23 can compensate for absence of IL-6 to promote SFB-directed Th17 cell differentiation. These results indicate that redundant cytokine signals guide commensal microbe-dependent Th17 cell differentiation in the MLN and accumulation of the cells in the lamina propria.

## Introduction

CD4^+^ T helper cells play crucial roles in vertebrate adaptive immune responses against potentially pathogenic microbes. CD4^+^ T cells additionally respond to antigens encoded by commensal microbes, and thereby contribute to the diversity of the T cell antigen receptor (TCR) repertoire resident within tissues. Differentiation of T cells into effector cells with diverse programs requires their activation through the TCR and CD28, the co-stimulatory receptor, as well as signaling through a multitude of cytokine receptors. Cytokines in the microenvironment, produced in large part by myeloid cells, direct the different programs of differentiation, which are readily distinguished by the induction of subset-specific transcription factors. Thus, interferon-γ (IFNγ)-producing Th1 cells express Tbet (*Tbx21*) (Szabo et al., 2002), Th2 cells that secrete interleukin 4 (IL-4), IL-5, and IL-13 express GATA3 (Agnello et al., 2003), and Th17 cells characterized by the production of IL-17A, IL-17F, and IL-22 express RORγt^+^ (Ivanov et al., 2006). In addition, induced regulatory T cells (iTreg) upregulate Foxp3 (and often other lineage-defining transcription factors, e.g. RORγt) and can produce IL-10 (Sefik et al., 2015) (Ohnmacht et al., 2015) (Xu et al., 2018).

Th17 cells are critical for epithelial homeostasis and host defense against bacteria and fungi at diverse mucosal sites (Korn et al., 2009). Dysregulated Th17 cell responses, however, promote several chronic inflammatory and autoimmune diseases such as inflammatory bowel disease (IBD), psoriasis, and various forms of arthritis (Maddur et al., 2012). Therefore, manipulation of Th17 cell responses has the potential to provide effective therapies for multiple immune system-based diseases. Developing therapies that do not hinder host-protective functions requires a thorough understanding of the molecular and cellular mechanisms underlying Th17 cell regulation during both homeostasis and chronic inflammation. In the past decade, the cytokine requirements for Th17 cell differentiation have been investigated *in vitro* and *in vivo* (Korn et al., 2009). *In vitro* differentiation of mouse Th17 cells from naïve CD4^+^ T cells can be achieved by culturing TCR/CD28-stimulated T cells with either TGF-β and IL-6 (Bettelli et al., 2006) (Mangan et al., 2006) (Veldhoen et al., 2006), with IL-6, IL-1β, and IL-23 (Ghoreschi et al., 2010), or with IL-6 and serum amyloid A (SAA) proteins (Lee et al., 2020). These conditions induce the expression of IL-23R and RORγt in a STAT3-dependent manner (Zhou et al., 2007). Th17 cell differentiation from naïve human cord blood CD4^+^ T cells can similarly be achieved with a combination of TGF-β, IL-1β, and IL-6, IL-21, or IL-23 (Manel et al., 2008). These cytokines have also been reported to be involved in pathogenic Th17 cell functions in the context of inflammatory conditions such as experimental autoimmune encephalomyelitis (EAE), mouse models of colitis, and IBD in humans (Bettelli et al., 2006) (Korn et al., 2007) (Langrish et al., 2005) (Nurieva et al., 2007) (Kamimura et al., 2003) (Cua et al., 2003) (Sanchez-Munoz et al., 2008) (Perrier and Rutgeerts, 2011) (Lee et al., 2020). IL-23, in particular, is critical for conferring pathogenicity in several animal models, but the transcriptional and metabolic features that distinguish homeostatic from pathogenic Th17 cells remain poorly understood.

The vertebrate gastrointestinal (GI) tract is colonized by hundreds of species of bacteria, which outnumber total host cells. Commensal microbes contribute not only to the activation of innate immune responses mediated by way of multiple sensing pathways, but also to the establishment of the peripheral adaptive immune repertoire and the acquisition of diverse T lymphocyte effector functions (Hooper et al., 2012). The generation of gut Th17 cells is especially dependent on commensal bacteria (Honda and Littman, 2016). While germ-free mice have few, if any, Th17 cells in their gut lamina propria (LP), specific pathogen-free (SPF) mice, which harbor many commensal species, have abundant intestinal Th17 cells that help maintain gut homeostasis (Ivanov et al., 2008) (Atarashi et al., 2008) (Hall et al., 2008). We previously showed that *Il-6* deficient mice have fewer IL-17A-producing CD4^+^ T cells in the intestine compared to *Il-6* sufficient mice (Ivanov et al., 2006). However, it was also reported that IL-6 is dispensable and, instead, microbiota-induced IL-1β is essential for the development of steady-state Th17 cells in the murine intestinal LP (Shaw et al., 2012). These studies did not account for immune responses that can vary according to which bacteria colonize the host. It is known that mice from different animal facilities and vendors have different intestinal microbiota (Chudnovskiy et al., 2016). The diversity of commensal communities in different vivaria likely accounts for conflicting results in inflammation-related studies, and may explain the different reported cytokine requirements for Th17 cell differentiation *in vivo*. We previously demonstrated that mono-colonization of germ-free mice with commensal segmented filamentous bacteria (SFB) is sufficient to induce Th17 cells that are specific for SFB antigens (Ivanov et al., 2009) (Yang et al., 2014). Recently, we described a two-step process for the functional differentiation of homeostatic Th17 cells in response to SFB colonization in the gut (Sano et al., 2015). In the first step, SFB antigen-specific CD4^+^ T cells are primed and polarized such that they are in a poised state, marked by the expression of RORγt in Th17 cells in the draining lymph nodes. In the second step, T cells undergo functional maturation in the intestinal LP, with triggering of Th17 signature cytokine gene expression programs by epithelial cell-derived factors. SAA1 and SAA2, produced by the epithelial cells in the ileum in response to SFB colonization, were shown to act on the poised RORγt^+^ Th17 cells, resulting in production of the effector cytokines IL-17A and IL-17F (Sano et al., 2015). In the current study, we have investigated the roles of cytokines and inductive sites in the discrete steps of homeostatic Th17 cell differentiation.

To explore the molecular mechanisms of Th17 cell induction, especially the expression of RORγt in vivo, we adoptively transferred SFB-specific TCR transgenic naïve mouse T cells into SFB-colonized hosts, allowing us to track SFB-specific Th17 cell expansion and differentiation. Here, we focused on characterizing the mechanisms by which SFB induces Th17 cells in the intestine-draining mesenteric lymph nodes (MLN) and the small intestine lamina propria (SILP). We found that SFB-induced expression of RORγt and proliferation of naïve T cells occurred in the MLN, but not in Peyer’s patches (PP), another inductive site in the intestinal mucosa. The RORγt^+^ Th17 cells migrated from the MLN to the SILP and PPs in an integrin β7-dependent manner. Although IL-6 was required for early RORγt expression by SFB-specific T cells in the MLN and SILP, IL-21 or IL-23, but not IL-1β, compensated in its absence at later times after bacterial colonization. These results indicate that Th17 cell differentiation occurs in discrete steps at different locations, and that several back-up systems exist to insure induction of the RORγt-directed program.

## Results

### Sites and kinetics of SFB-specific Th17 cell induction and proliferation

To explore the molecular and cellular mechanisms of Th17 cell induction *in vivo*, we first tracked SFB-induced Th17 cell expansion and differentiation based on cell division and RORγt expression, respectively. To monitor when, where, and how naïve T cells differentiate into Th17 cells upon SFB colonization, we adoptively transferred carboxyfluorescein succinimidyl ester (CFSE)-labeled naïve T cells from SFB-specific TCR transgenic (7B8) mice into SFB-gavaged host mice (Yang et al., 2014). Two days following the transfer of 50,000 naïve 7B8 T cells, some of the transferred cells began to express CD25 (data not shown) and RORγt in the intestine-draining mesenteric lymph nodes (MLN). Notably, at this time point, there was no proliferation of these cells, and CD25 (data not shown) and RORγt^+^ 7B8 T cells were undetectable in the Ileum, PP, or spleen (Fig.1A). By day 3, 7B8 T cells within the MLN had undergone several rounds of proliferation and many expressed RORγt (Fig.1A) but not Foxp3 (data not shown). Donor-derived 7B8 T cells only began to accumulate in the ileal LP at 4 or 5 days following T cell transfer (Fig.1A and B). By 7 days post-transfer, most 7B8 T cells in the intestinal LP expressed RORγt (Fig.1A). We also checked other tissues to determine where SFB-specific Th17 cells differentiate. We found some proliferating RORγt^+^ 7B8 T cells in the spleen and Peyer’s patches (PP) before we observed RORγt^+^ 7B8 T cells in the intestinal LP (Fig.1A). Overall, expanded 7B8 T cells were detected in the MLN before any of the other tissues tested (Fig. 1A and 1B).

**Figure 1.**
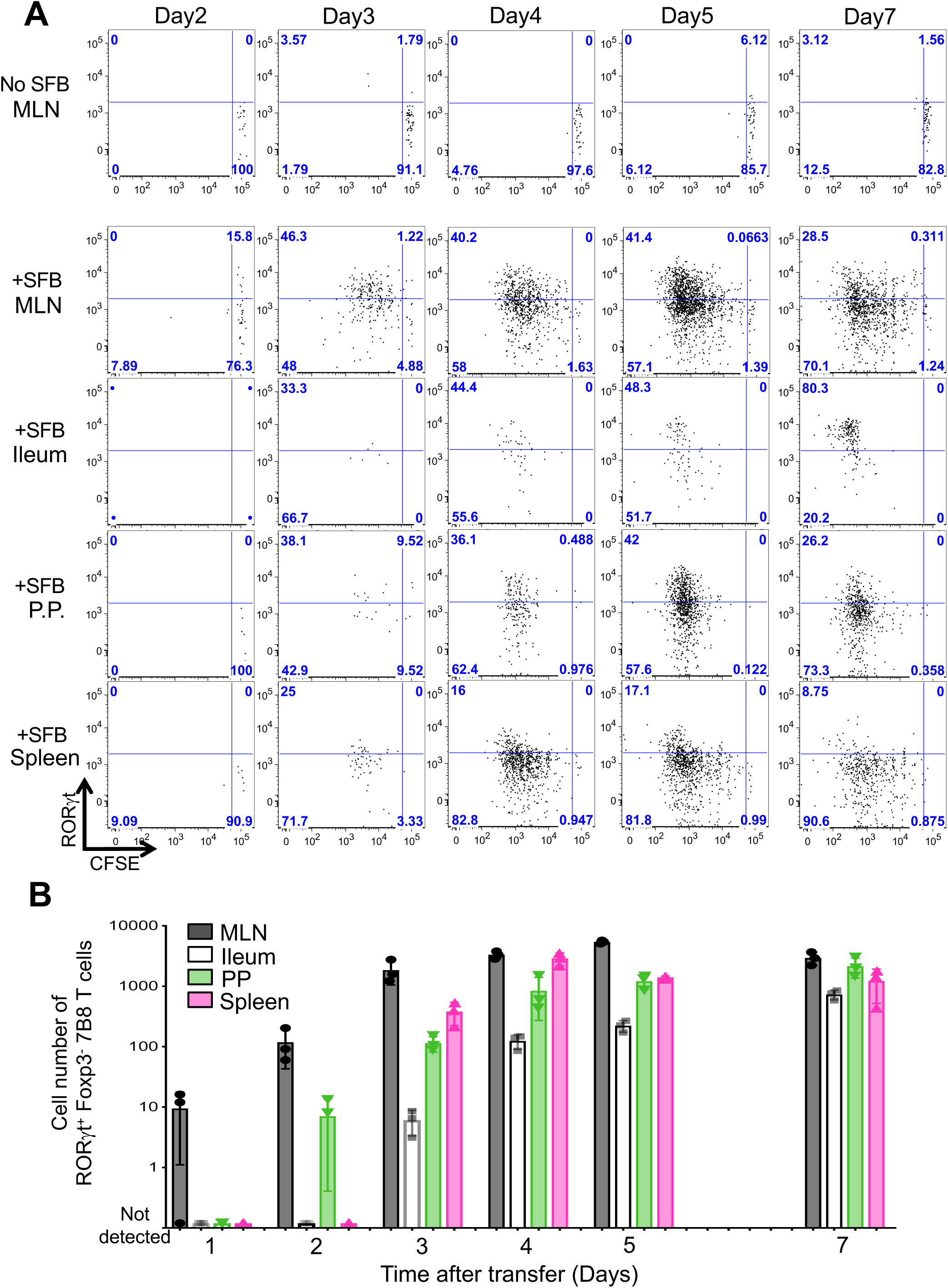
Kinetics of SFB-specific Th17 cell differentiation in tissues of SFB-colonized mice. **(A)** Cell expansion, monitored by CFSE dilution, and RORγt expression in donor-derived 7B8 T cells from diverse tissues of SFB-gavaged mice. 50,000 naïve 7B8 TCR transgenic T cells (*Cd45.1/Cd45.2*) were transferred into SFB gavaged mice (*Cd45.2/ Cd45.2)*. SFB-specific 7B8 T cells were defined as CD4^+^, TCRβ^+^, TCRVβ14^+^, and CD45.1^+^ cells. As a negative control, SFB-free SPF B6 mice were used. The time course experiment was performed once, with 3 mice per time point. Repeats with specific time points (Day2, Day3, Day4, Day5, and Day7) were performed multiple times using biological duplicates, with similar results. **(B)** The number of SFB-specific Th17 cells in the MLN, Ileum, PPs, and spleens of SFB-gavaged C57BL/6 mice. SFB-specific Th17 cells were defined as Foxp3^-^ RORγt^+^ 7B8 T cells described in Figure 1A. These data were generated from one time course experiment and represent the mean from 3 or 4 mice +/- SD. See also Figure **S1**.

To test our model under conditions that better match the physiological frequency of antigen-specific T cells within a host, we transferred only 5000 naïve 7B8 T cells into SFB-gavaged mice. We again first detected Th17 cell expansion and differentiation in the MLNs and, at later times, in the intestinal LP (Fig. S1A and S1B), suggesting that they had migrated from sites of induction. SFB colonize and adhere to epithelial cells in the terminal ileum (Koopman et al., 1987), but not in the duodenum and colon (Sano et al., 2015). Nevertheless, we observed similar numbers of RORγt^+^ 7B8 T cells in the duodenum and colon when compared to the ileum (Fig.S1A).

It has been proposed that PPs are the primary site of SFB-specific Th17 cell induction (Lecuyer et al., 2014). In addition, PP dendritic cells were shown to endow CD8^+^ T cells with small intestine-homing capability, through induction of α4β7 and CCR9 (Mora et al., 2003). We therefore compared PP and MLN tissue sections to identify clusters of SFB-specific Th17 cells at different times after naïve 7B8 T cell transfer. In the absence of SFB, there were no donor-derived RORγt^+^ Th17 cells in either MLN or PPs (data not shown). In SFB-gavaged mice, however, we were able to detect clusters of SFB-specific T cells in the MLNs and PPs. Many of these clustered cells expressed RORγt (Fig. 2A) and stained positive for the proliferation marker Ki67 (Fig. S2A), suggesting that these were SFB-specific Th17 cells undergoing active differentiation. Notably, these clusters were detected only in MLNs at 2 and 3 days following transfer, and only on day 4 were some clusters detected in PPs (Fig. 2B).

**Figure 2.**
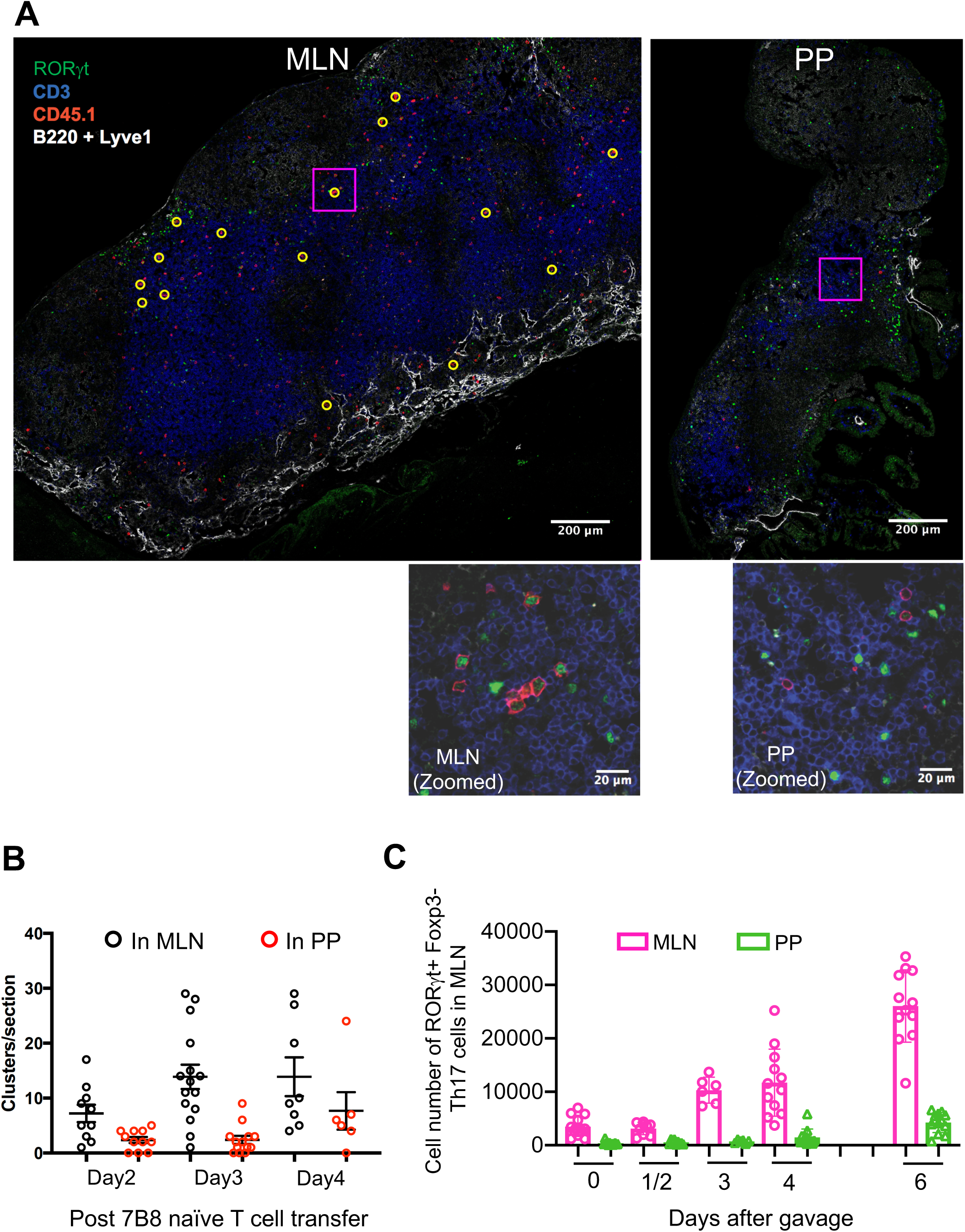
Quantification of Th17 cells in the MLN and PPs following SFB gavage. **(A)** Th17 cell clusters in MLN and PPs 3 days following transfer of 50,000 naïve 7B8 T cells. Sections were stained with anti-RORγt, anti-CD3, anti-CD45.1, anti-B220, and anti-Lyve-1 and were analyzed by confocal microscopy. B220 and Lyve1 were both stained with Pacific Blue-conjugated antibodies, but can be distinguished by cell morphology and stain intensity. Yellow circles indicate clusters of CD45.1^+^ cells, defined as 2 or more CD45.1^+^ cells in contact. Representative of 5 mice from 2 experiments. Insets are enlargements of the boxed areas. **(B)** Quantification of clustering in MLN and PPs imaged as in (A). Numbers of clusters per tissue section are shown. Representative of two experiments, each with 2-3 mice per time point. Each symbol represents a single tissue slice, and the bar indicates average +/- SD. **(C)** Kinetics of Th17 cell accumulation in the MLN and PPs following SFB gavage. The cell numbers were calculated based on total cells in the MLN and PPs. Experiments were repeated twice with similar results and results were all combined (SFB gavaged mice, Day0: n=13, Day1/2: n=9 (Day1: n=3, Day2: n=6), Day3: n=6, Day4: n=12, Day6: n=12). Data are represented as mean +/- SD. See also Figure **S2**.

We also quantified the number of endogenous Th17 cells in the MLN and PP of mice following gavage with SFB. We found that while the number of RORγt+ Th17 cells began to increase in PPs at 6-days post SFB gavage, those in the MLN began to expand within 3 days following SFB inoculation (Fig. 2C). To evaluate whether APCs from MLN carry SFB antigens, we collected APCs from MLN following SFB gavage and co-cultured with SFB-specific naïve T cells *in vitro*. Naïve 7B8 T cells expressed the early T cell activation markers CD25, CD69, and CD62L when they were co-cultured with APCs isolated from MLN at least 4 days after gavage, but not with APCs from ungavaged mice (Fig. S2B and S2C).

We next investigated whether Th17 cells can accumulate in the intestinal LP in response to SFB gavage if the mice lack PPs. PP development was abrogated by injection of LTβR-Ig fusion protein into dams at E16.5 and E18.5 of fetal development (Rennert et al., 1996). Although PPs were absent in treated mice (Fig. S2D), there was no effect on the number of SFB-specific Th17 cells in the mouse gut at 7 and 10 days following SFB gavage (Fig. S2E and S2F). Importantly, MLNs were intact and had similar numbers of RORγt^+^ Th17 cells in mice depleted of PPs and control antibody-injected mice (data not shown). These results indicate that PPs are dispensable for the induction of Th17 cells and for their expansion in the SILP in response to SFB gavage.

### Integrin β7 is essential for the migration of Th17 cells from the MLN to the intestine

Integrins α4 and β7 are induced on T cells in gut-draining lymph nodes and have a key role in homing of the T cells to intestinal LP (Schweighoffer et al., 1993) (Erle et al., 1994) (Johansson-Lindbom and Agace, 2007). Indeed, naïve 7B8 T cells transferred into SFB-colonized mice upregulated the expression of integrin α4 and β7 heterodimers on their surface following proliferation in response to antigen (Fig. 3A). To confirm that Th17 cells migrate to the intestine after expansion in MLN (Fig. 1 and 2), we co-transferred naïve CFSE-labeled *Itgb7*^+/-^ (*Cd45.1/ Cd45.2*) and *Itgb7*^-/-^ (*Cd45.2/Cd45.2*) 7B8 T cells into SFB-colonized hosts (*Cd45.1/Cd45.1*), and tracked the isotype-marked cells in various tissues at several time-points. Both *Itgb7*^*-/-*^ 7B8 and *Itgb7*^*+/-*^ 7B8 T cells expanded in the MLN and spleen at 4-days post co-transfer (Fig. 3B and 3C). However, *Itgb7*^*-/-*^ 7B8 T cells did not accumulate in the intestinal LP compared to *Itgb7*^*+/-*^ 7B8 T cells at 4, 7, and 10 days after transfer, but had increased accumulation in spleen (Fig. 3B, 3C, S3A and S3B). Taken together, we conclude that initial expansion of Th17 cells occurs primarily in the MLN, and SFB-specific Th17 cells subsequently migrate into the intestinal LP and PPs in an integrin β7-dependent manner. These data thus imply that APCs present SFB antigens and prime and polarize Th17 cells in the MLN.

**Figure 3.**
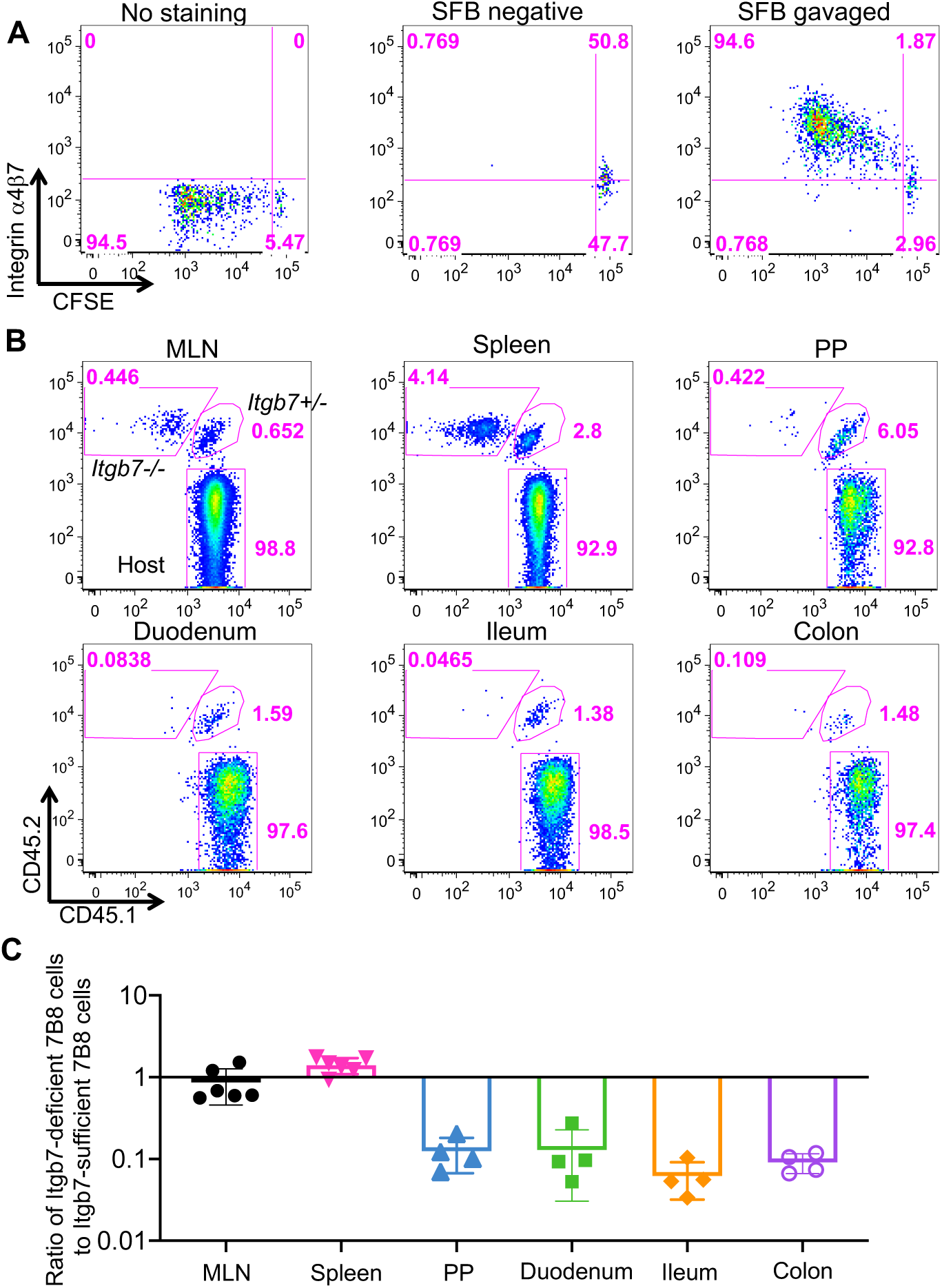
The role of α4β7 integrin in migration of SFB-specific Th17 cells from MLN to local tissues. **(A)** Expression of α4β7 integrin heterodimer in primed SFB-specific T cells in the MLN. 50,000 CFSE-labeled naïve T cells from SFB-specific TCR transgenic (7B8) mice were adoptively transferred into SFB-colonized mice. No staining represents the sample with isotype control IgG. Similar results were observed in multiple mice (n=4) at day 4 post adaptive transfer. **(B)** Localization of integrin β7-sufficient and -deficient SFB-specific T cells following adoptive transfer. 50,000 naïve T cells from *Itgb7* heterozygous (*Cd45.1/Cd45.2*) and homozygous mutant (*Cd45.2/Cd45.2*) 7B8 mice were co-transferred into SFB-gavaged C57BL/6 (*Cd45.1/Cd45.1*) mice. Four days following transfer, isotype-marked 7B8 T cells from multiple tissues were analyzed. **(C)** Ratios of *Itgb7-*deficient to *Itgb7*-sufficient (heterozygous) 7B8 T cells from multiple tissues in the same host mice at 4 days post transfer. Each symbol represents a single mouse, and the color bar indicates average +/- SD. Experiments were repeated twice and results were combined. n= 4 (Duodenum, Ileum, and colon); n= 6 (MLN and PPs). See also Figure **S3**.

### Cytokine requirements for SFB-induced Th17 cell differentiation *in vivo*

Antigen presentation is the initial step in the priming of antigen-specific T cells. Simultaneously, innate immune cell-derived cytokines direct differentiation of the T cells towards diverse effector programs marked by expression of the signature transcription factors. Thus, most of the 7B8 T cells located in the ileum in SFB-colonized mice expressed RORγt (Sano et al., 2015), but proliferating 7B8 T cells in the spleen, which may have migrated from the MLN, expressed little or no RORγt (Fig. S1C). We hence hypothesized that APCs in the MLN provide Th17-inducing cytokines, but continued RORγt expression in differentiated Th17 cells may require maintenance by cytokines in tissue microenvironments. Therefore, we next focused on cytokine requirements for Th17 cells in the MLN and intestinal LP. *In vitro* studies revealed that the combination of several cytokines including TGF-β, IL-1β, IL-6, IL-21, and IL-23 can induce RORγt expression and IL-17A production in CD4^+^ T cells (Acosta-Rodriguez et al., 2007; Bettelli et al., 2006; Manel et al., 2008; Mangan et al., 2006; Veldhoen et al., 2006; Wilson et al., 2007; Yang et al., 2007; Zhou et al., 2007). However, the identity of cytokines involved in RORγt expression *in vivo* remains unresolved. For example, whereas we described a requirement for IL-6 in the differentiation of intestinal CD4^+^ T cells that express RORγt, IL-23R, IL-17A and IL-17F (Ivanov et al., 2006), Nunez and colleagues demonstrated that IL-1β, but not IL-6, was required for steady state Th17 cell differentiation in gut (Shaw et al., 2012). To elucidate which cytokines are important for RORγt expression in SFB-induced Th17 cells, we orally introduced SFB-enriched fecal contents into SFB-free *Il-21r, Il-23 p19*, and *Il1r1* deficient mice. Seven or 10-days post SFB gavage, the numbers and proportions of RORγt^+^ Th17 cells in the ileum of these cytokine signaling-deficient mice were similar to those of littermate control mice (Fig. S4). In contrast, there were markedly fewer RORγt^+^ Th17 cells in the ileum of SFB-colonized *Il-6*^-/-^ mice compared to IL-6-sufficient littermate controls (Fig. 4A and 4B). Although this defect was pronounced during the first week after SFB gavage, there was a gradual increase in ileal RORγt^+^ Th17 cells in *Il-6*^*-/-*^ mice during the subsequent two weeks (Fig. 4A and 4B).

**Figure 4.**
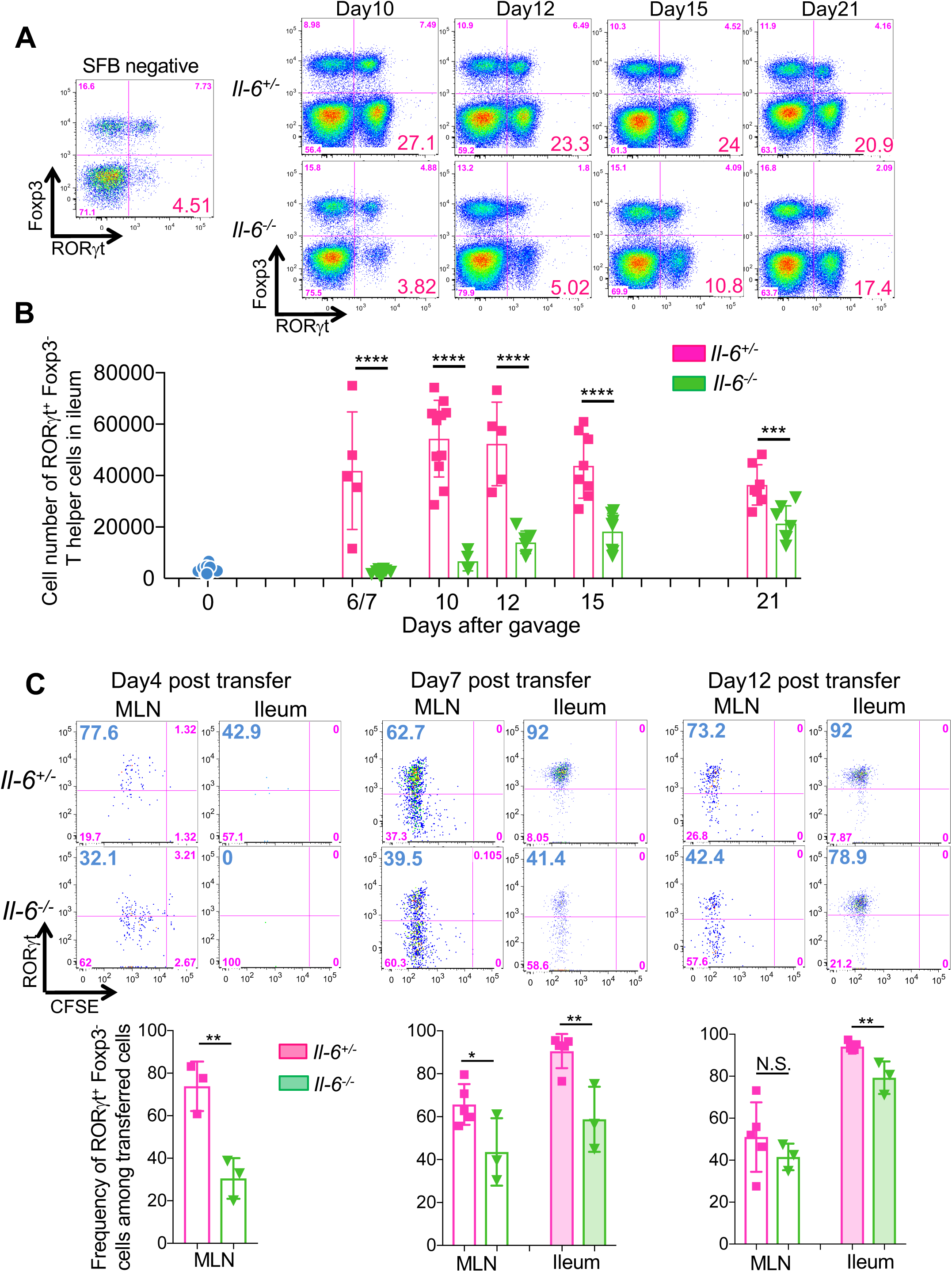
Early requirement for IL-6 in SFB-induced Th17 cell differentiation. **(A and B)** IL-6 requirement for Th17 cell differentiation in response to SFB gavage. IL-6-sufficient (heterozygous) and -deficient mice were gavaged with SFB and RORγt^+^ Foxp3^-^ Th17 cells in the ileal LP were assessed by FACS at different times. Representative FACS panels (A) and cell numbers (B). Results were combined from two experiments with littermates. Dots indicate individual mice and the color bars indicate mean values +/- SD. **(C)** Requirement for IL-6 in SFB-specific Th17 cell differentiation in the MLN and ileum. 50,000 (for Day 4) or 5,000 (for Day 7 and Day 12) naïve T cells from SFB Tg (7B8) mice (*Cd45.1/Cd45.2*) were adoptively transferred into SFB-gavaged *Il-6* sufficient and deficient mice (*Cd45.2/Cd45.2*). Donor-derived T cells in the MLN and ileum were analyzed for RORγt expression on the indicated days. Representative FACS profiles in top panels, aggregate data in bottom panels. Each symbol represents a single mouse, and the color bar indicates average +/- SD. *<0.05, **<0.001, *** p<0.001, and **** p<0.0001. See also Figure **S4** and **S5**.

To confirm these results with SFB-induced antigen-specific T cells, we transferred CFSE-labeled 7B8 naïve T cells into SFB-gavaged *Il-6*^*+/-*^ *and Il-6*^*-/-*^ mice. Four days following transfer, proliferation of 7B8 T cells in MLN of *Il-6*^*-/-*^ mice was similar to that in *Il-6*^*+/-*^ mice based on CFSE dilution (Fig. 4C). However, there was substantial reduction of RORγt-expressing 7B8 T cells in the MLN of *Il-6*^*-/-*^ compared to *Il-6*^*+/-*^ mice (Fig.4C). Similarly to the T cells in the MLN, there were fewer RORγt-expressing SFB-specific T cells in the ileum of *Il-6*^*-/-*^ hosts at 7 days following transfer (Fig. 4C). However, expression of RORγt in transferred 7B8 T cells was similar in *Il-6*^*-/-*^ and *Il-6*^*+/-*^ mice at the later time points (Fig. 4C). To confirm IL-6 dependency using different mouse models, we also quantified the number of Th17 cells in *Il-6*^*+/-*^ and *Il-6*^-/-^ mice that had been stably colonized with SFB and various other commensals in a SPF room of the animal facility at the New York University Skirball Institute (NYU^SK^). Long-term SFB-colonized *Il-6*^-/-^ mice in the NYU^SK^ facility had a moderate, but statistically significant, reduction in Th17 cells compared to littermate *Il-6*^*+/-*^ mice (Fig. S5A and S5B). We also tested other microbial communities that can induce Th17 cells in the mouse gut. Mice from Jackson Laboratory (JAX), which have a microbiota lacking SFB, consistently contain a small population of intestinal RORγt^+^ Th17 cells (Ivanov et al., 2008). We quantified the number of RORγt^+^ Th17 cells in IL-6-sufficient and -deficient littermates bearing JAX flora, and found a similar reduction in RORγt^+^ Th17 cells in the MLN and ileum of the homozygous mutant mice (Fig. S5C and S5D). Taken together, these results indicate that, following colonization with Th17-inducing bacteria, IL-6 is a strong early inducer of RORγt expression in responsive T cells in the MLN. However, it is not absolutely required for Th17 cell differentiation, suggesting that there are other cytokines capable of inducing RORγt in intestinal T cells.

### Stat3 is required for Th17 cell differentiation in response to SFB colonization

Because at least some RORγt^+^ Th17 cell differentiation proceeds in the absence of IL-6 (Fig. 4 and S5), we speculated that other cytokines may compensate. Multiple cytokines, including IL-1β, IL-6, IL-21, and IL-23, have been reported to induce RORγt expression during *in vitro* Th17 cell differentiation, and several of these function through stimulation of Stat3 phosphorylation. Moreover, it was reported that Stat3 signaling is required in IL-17-producing Th17 cells in multiple tissues including MLN and intestinal LP (Harris et al., 2007). To investigate whether Stat3-stimulating cytokines compensate for loss of IL-6 in microbiota-dependent induction of RORγt expression in CD4^+^ T cells *in vivo*, we next examined whether SFB-dependent Th17 cell differentiation can occur in the absence of Stat3. We quantified RORγt-expressing Th17 cells in mice that were stably colonized with SFB and lacked expression of *Stat3* in T cells (*Cd4*^*cre/+*^ *Stat3*^*flox/flox*^). As previously reported (Harris et al., 2007) (Ohnmacht et al., 2015), *Stat3* deficiency in CD4^+^ T cells led to a substantial reduction in both frequency and numbers of RORγt^+^ Th17 cells in the MLN, to almost complete absence of such cells in the ileum, and to loss of RORγt^+^ induced Treg cells (Fig. S6A and S6B). The reduction in Th17 cells was much more marked, particularly in the ileum, in the absence of Stat3 as compared to the absence of IL-6. This result suggested that Stat3-stimulating cytokines other than IL-6, particularly IL-21 and IL-23, may contribute to RORγt expression in Th17 cell differentiation *in vivo*.

### IL-21 and IL-23, but not IL-1βγ compensate for the lack of IL-6 in SFB-dependent Th17 cell differentiation

To determine which cytokines contribute to Th17 cell differentiation in the absence of IL-6 *in vivo*, we prepared isotype-marked 7B8 TCR transgenic mice deficient in *Il-1r1, Il-21r*, or *Il-23r*. We first determined whether IL-23R is expressed on 7B8 T cells in the *Il-6*^*-/-*^ host colonized with SFB. Indeed, there was reduced GFP expression in the SFB-specific T cells from *Il-23r*^*gfp/+*^ reporter mice in *Il-6*^*-/-*^ compared to *Il-6*^*+/-*^ recipient mice, consistent with partial dependency of IL-23R expression on IL-6 (Zhou et al., 2007) (Fig. S6C). We then co-transferred *Il-23r*^*gfp/+*^ (*Cd45.1/Cd45.1*) and *Il-23r*^*gfp/gfp*^ (*Cd45.1/Cd45.2*) naïve 7B8 T cells into stably SFB-colonized *Il-6*^*+/-*^ and *Il-6*^*-/-*^ hosts (*Cd45.2/Cd45.2*). In *Il-6*^*+/-*^ hosts, there was little difference in RORγt^+^ induction between *Il-23r*^*gfp/+*^ and *Il-23r*^*gfp/gfp*^ 7B8 T cells in the MLN and the ileum. In *Il-6*^*-/-*^ hosts, however, there was substantial reduction in RORγt expression among *Il-23r*-deficient 7B8 T cells compared to *Il-23r* -sufficient cells in MLN and the ileum (Fig. 5A and 5B). Transferred *Il-21r*^*-/-*^ 7B8 T cells similarly failed to express RORγt in the MLN and ileum in *Il-6*^*-/-*^ hosts but differentiated normally into Th17 cells in *Il-6*-sufficient mice (Fig. 5C and 5D). Notably, *Il-1r1* deficiency did not affect RORγt expression even in the absence of IL-6 signaling *in vivo* (Fig. S6D and S6E).

**Figure 5.**
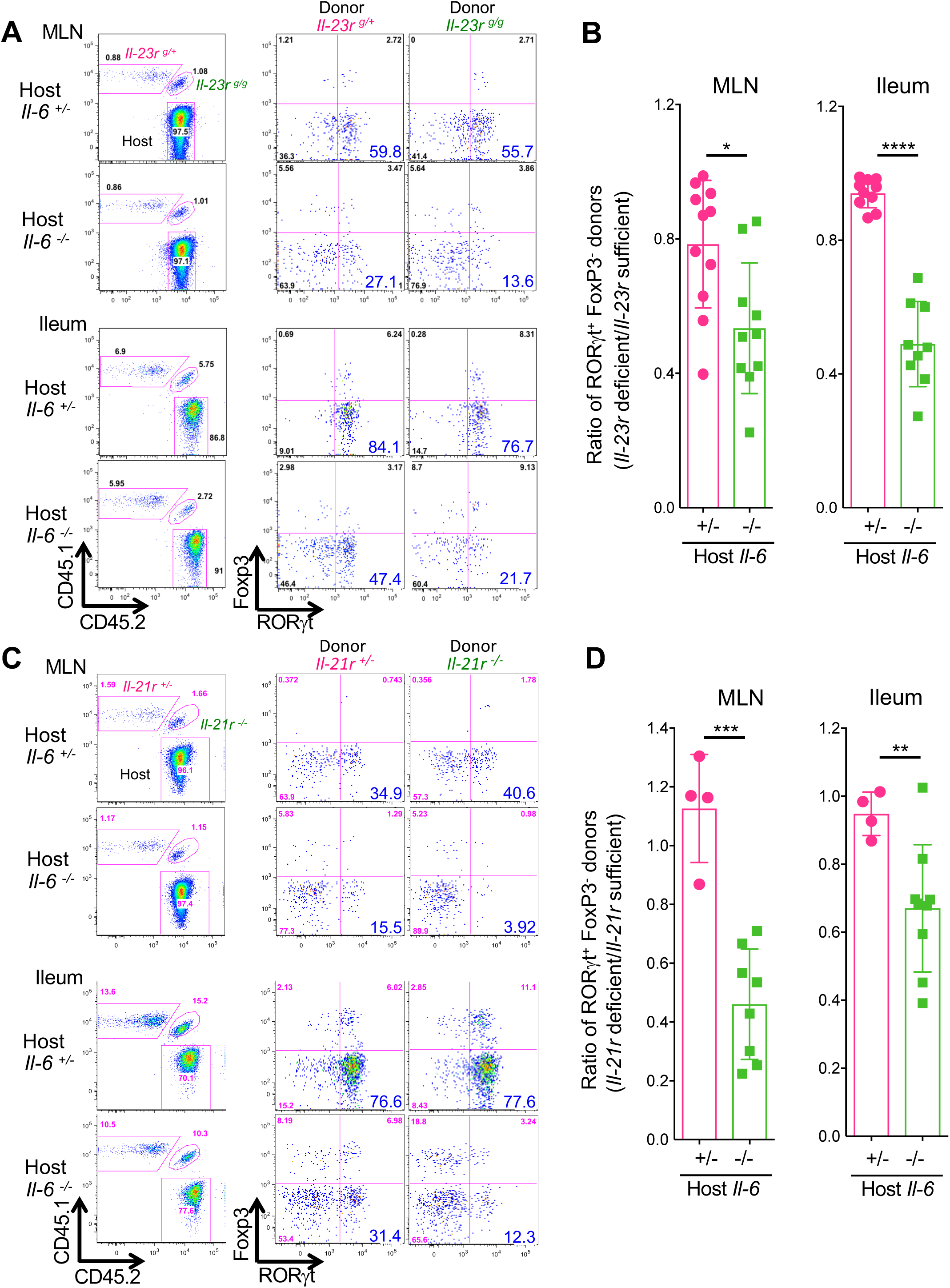
Cytokine redundancy in the differentiation of SFB-specific Th17 cells in the MLN and ileum. **(A-D)** Requirement of IL-23R (A and B) and IL-21R (C and D) signaling for SFB-specific Th17 cell differentiation in *Il-6*-sufficient and -deficient mice. 5,000 naïve T cells from cytokine receptor sufficient and deficient isotype-marked 7B8 TCR transgenic mice were co-transferred into SFB-gavaged *Il-6* sufficient and deficient mice (*Cd45.2/Cd45.2*). Representative FACS plots of RORγt and Foxp3 expression in ileal 7B8 T cells with homozygous or heterozygous *Il-23r* and *Il-21r* mutations at 7 days post transfer into *Il-6*-sufficient and -deficient mice (A and C). Panels at left indicate gating for donor-derived versus host T cells. Ratios of RORγt^+^ Foxp3^-^ Th17 cells (cytokine receptor sufficient vs deficient T cells) in *Il-6* sufficient and deficient mice (B and D). Experiments were performed twice with littermate controls. All results were combined. Dots indicate individual mice and color bars indicate average +/- SD. For IL-23R signaling (*Il6*^*+/-*^: n=9, *Il6*^*-/-*^: n=10). For IL-21R signaling (*Il6*^*+/-*^: n=4, *Il6*^*-/-*^: n=9). * p<0.05, ** p<0.01, *** p<0.001, and **** p<0.0001. See also Figure **S6**.

We next compared Th17 cell differentiation in mice deficient for different combinations of the Stat3-signaling cytokines. In IL-6, IL-21R, and IL-23R triple-deficient mice, RORγt-expressing Th17 were barely detectable in the ileum of SFB-colonized mice at steady state (Fig 6A and B). These animals were housed in the vivarium at the University of Illinois at Chicago (UIC), and control, single-, or double-deficient mice recapitulated the partial loss of Th17 cells observed in the NYU^SK^ mice. The number of RORγt-expressing Th17 cells in the ileum of IL-21R/IL-23R double-deficient mice was similar to that in the sufficient mice (Fig 6A and B), and deficiency of IL-6 was required to observe defective Th17 cell differentiation. The results thus indicate that signaling through receptors for IL-21 and IL-23, but not IL-1β, can partially compensate for the loss of IL-6 in differentiation of SFB-specific Th17 cells in the mouse intestine.

**Figure 6.**
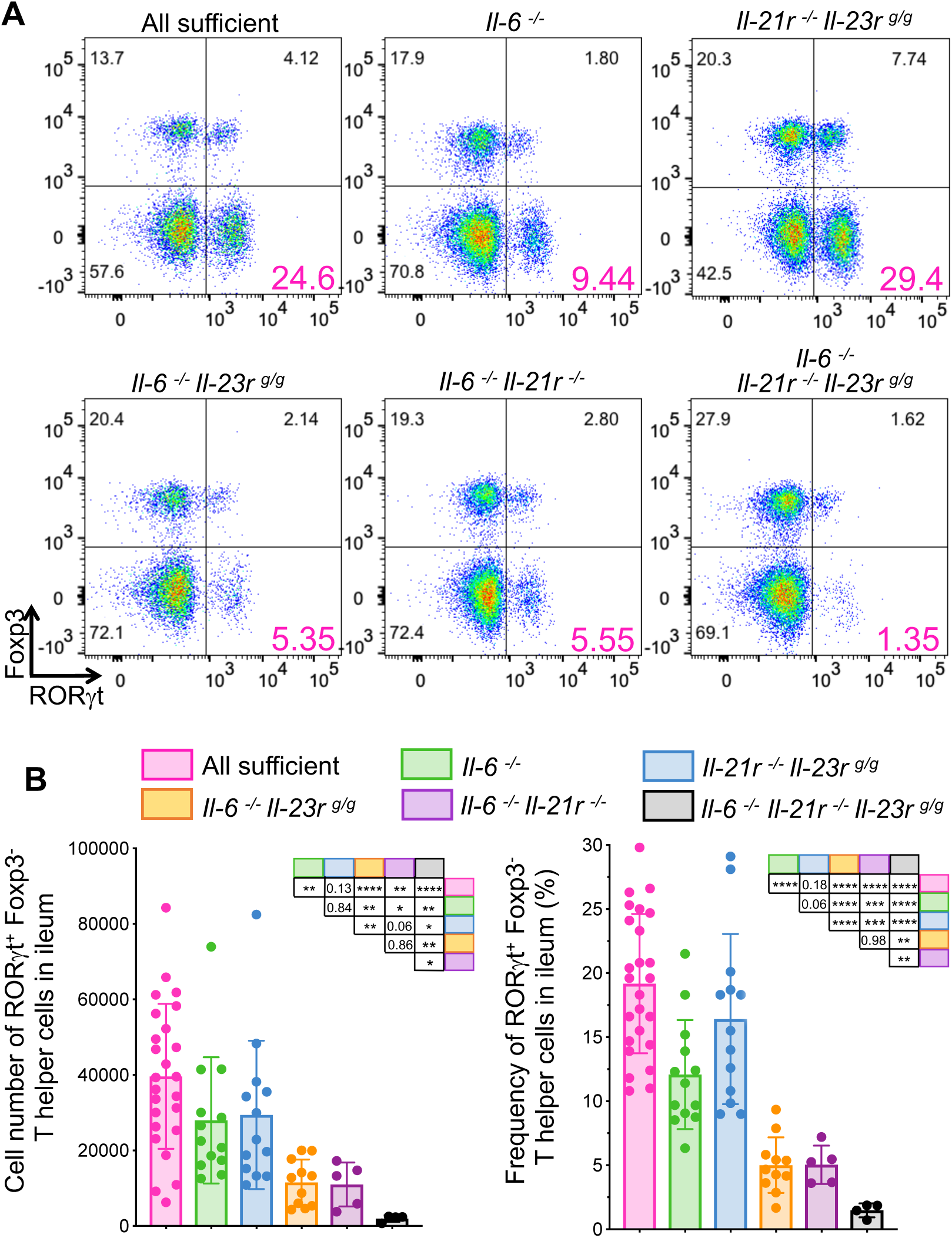
Contribution of multiple cytokines to Stat3-dependent signaling redundancy in SFB-induced Th17 cell differentiation. **(A and B)** Representative FACS plots of ileal Th17 cells (RORγt^+^Foxp3^-^) in stably SFB-colonized mice with mutations in cytokines or their receptors (A). Cell numbers (B, left) and frequencies (B, right) of Th17 cells in the ileum. Experiments were repeated with indicated genotypes at least three times and results are combined. Littermates were used in most cases. Mice were confirmed as SFB^+^ by qPCR or microscopic observation. *p*-values for differences between each group of mice are shown in the tables. Dots represent individual mice and color bars indicate average +/- SD. * p<0.05, ** p<0.01, *** p<0.001, and **** p<0.0001.

## Discussion

We showed in this report that Th17 cell differentiation in response to colonization with the commensal microbe SFB is primarily initiated in the gut-draining MLN and that the requisite T cell intrinsic Stat3 signaling can be achieved through the action of cytokines that can compensate in part for each other’s absence. We found that IL-6 is a potent early inducer of RORγt in SFB-specific Th17 cells in the draining lymph nodes, but, in its absence, IL-21 and IL-23 can compensate to activate the Th17 program. Our study also shows that, after initial differentiation in the MLN, SFB-specific Th17 cells migrate to the intestinal LP in an integrin β7-dependent manner. Finally, RORγt^+^ Th17 cells that have migrated from the MLN accumulate in the intestinal LP and are further stimulated by the tissue microenvironment to produce effector cytokines (Sano et al., 2015). RORγt^+^ Th17 cells localized at mucosal sites likely serve as a reservoir of microbiota-induced cells that can be reactivated in response to tissue damage and thus maintain homeostasis (Honda and Littman, 2016). However, Th17 cells also contribute to autoimmune disease and inflammatory pathology in humans, and IL-1β, IL-6, and IL-23 are required in Th17-dependent mouse models of autoimmunity (Kamimura et al., 2003) (Chen et al., 2006). Furthermore, antibodies targeting these cytokines or their receptors have been clinically effective for treating rheumatoid arthritis and inflammatory bowel disease in humans (Jones et al., 2011) (Gaffen et al., 2014) (Tanaka and Kishimoto, 2012). Our results suggest that such treatments may not fully block homeostatic Th17 cell differentiation, as IL-6 and IL-23 signaling are partially redundant for RORγt induction and Th17 cell differentiation.

### Microbiota-dependent Th17 cell priming and differentiation occurs in MLNs

It has been reported that SFB induces bacterial antigen-specific IgA and Th17 cell responses in secondary and tertiary lymphoid tissues, including Peyer’s patches and isolated lymphoid follicles (ILFs) (Lecuyer et al., 2014) (Teng et al., 2016). On the other hand, it has also been suggested that Th17 cells are generated even in mice lacking secondary lymphoid organs (Goto et al., 2014) (Geem et al., 2014). Here, we have demonstrated that the MLNs are the initial site for RORγt^+^ Th17 cell differentiation. SFB-specific RORγt^+^ Th17 cells were detected in similar numbers not only in SFB-colonized ileum but also in the lamina propria of duodenum and colon, which are not colonized with SFB (Sano et al., 2015). Moreover, RORγt^+^ Th17 cells in the intestinal LP were detected much later than those in the MLN and PPs. These results indicated that the intestinal LP and ILFs are not the predominant nor initial sites of SFB-specific Th17 cell differentiation. We therefore investigated the other candidate sites for Th17 cell priming and differentiation, the MLN and PPs. RORγt and CD25 were upregulated early in MLN, but not PPs, spleen or intestinal LP, after transfer of SFB-specific 7B8 cells. In addition, clusters of SFB-specific RORγt^+^ Th17 cells were observed in the MLN, but not in the PPs, at early time points. Moreover, even in mice depleted of PPs, RORγt^+^ Th17 cells were detected in the MLN and ileum LP with similar kinetics and abundance to control mice. The integrin α4β7, which mediates lymphocyte migration to intestinal lamina propria, was critical for accumulation of SFB-specific Th17 cells in the small intestine, but not for their induction in the MLN. Finally, APCs from MLN stimulated SFB-specific naïve T cells *in vitro* early after SFB colonization. Based on these results, we conclude that the MLN, rather than PPs or intestinal LP, is the principal site for early SFB-specific Th17 cell differentiation. However, our results do not rule out the possibility that Th17 cells can be also generated in the PPs and gut associated lymphoid tissue (GALT). Th17 cells were induced in the intestine of LTα-deficient mice, which lack secondary and tertiary lymphoid tissues, but the kinetics of induction were not compared to those in control mice (Goto et al., 2014). In the absence of secondary lymphoid tissues, APCs carrying SFB antigens may prime T cells locally in addition to maintaining homeostatic Th17 cells in the ileal LP. Under these non-physiological conditions, commensal-specific T cells may thus be primed ectopically.

CFSE-diluted 7B8 T cells were observed in spleen in addition to the MLNs and PPs. More integrin β7-deficient 7B8 cells accumulated in the spleen relative to integrin β7-sufficient 7B8 T cells. These results indicate that commensal-induced Th17 cells generated in the MLN have the potential to circulate systemically. RORγt^+^ Th17 cells home from the MLN to their target tissues based on their expression of distinct chemokine receptors and integrins, and failure to home to the intestinal LP likely results in more accumulation in blood and spleen. The 7B8 cells in spleen expressed reduced amounts of RORγt compared to the cells in MLN or intestine. This may reflect a requirement for a microenvironment in which there is continued antigen presentation or production of soluble factors, such as serum amyloid A proteins, to maintain expression of RORγt.

### Cytokine requirements and redundancy during commensal-induced Th17 cell differentiation

Studies on conditions required for *in vitro* Th17 cell differentiation have reported the importance of several cytokines, including IL-1β, IL-6, IL-21, and IL-23 (Korn et al., 2009). Here, we showed that IL-6 was required *in vivo* for early induction of RORγt expression in SFB-specific Th17 cells. However, in its absence, other Stat3-stimulating cytokines, IL-21 and IL-23, provided the necessary stimuli for the Th17cell differentiation program in mouse intestine. IL-21 is known to be produced by activated CD4^+^ T cells following IL-6 stimulation (Korn et al., 2007) (Zhou et al., 2007) (Nurieva et al., 2007). IL-6 and IL-21 induce IL-23R expression on activated T cells (Zhou et al., 2007). Induced RORγt itself can also enhance the expression of IL-23R on Th17 cells (Wang et al., 2015). These features may provide positive feedback and backup systems to maintain Th17 cells in the gastrointestinal tract.

Unlike IL-21 and IL-23, IL-1β failed to compensate for the loss of IL-6 in rescuing SFB-dependent Th17 cell differentiation. However, IL-1β has been shown to be required for pathogenic functions of Th17 cells, e.g. in EAE. In addition, IL-1β, but not IL-6, was reported to be required for microbiota-induced gut Th17 cell differentiation in mice that were colonized with SFB (Shaw et al., 2012). A parsimonious explanation for these disparate findings is that SFB induces a different set of cytokines in the context of the microbiota in different vivaria. The frequency of SILP-localized RORγt^+^ Th17 cells reported by Shaw et al. in IL-1R1 sufficient mice was considerably lower than what we observed, consistent with this possibility.

IL-1R1 signaling deficiency not only affects the differentiation of Th17 cells, but also impairs the function of RORγt^+^ Th17 cells in the healthy gut and in the spinal cord and LNs during EAE (Chung et al., 2009; Sha and Markovic-Plese, 2011; Shaw et al., 2012). Although IL-1β may stimulate the Stat3 pathway indirectly (Mori et al., 2011), IL-1R1 signaling may have different roles and redundant functions in homeostatic versus pathogenic Th17 cell differentiation. It was reported that T cell-intrinsic IL-1R1 signaling licenses effector cytokine production, stabilizing Th17 and Th2 cytokine transcripts (Jain et al., 2018). There may also be non-T cell-intrinsic functions of IL-1R1 that contribute to Th17 cell differentiation and that were not examined in our model.

### Microbiota-induced Th17 cells in autoimmune disease

The SFB colonization model that we and others study is considered to result in differentiation of non-pathogenic or homeostatic Th17 cells. However, under some circumstances, SFB can contribute to systemic autoimmune disease. Thus, SFB mono-colonized mice had more severe disease in the K/BxN autoimmune arthritis model (Wu et al., 2010), and IL-17A-producing SFB-specific Th17 cells were detected in the spleen and lung of K/BxN mice (Bradley et al., 2017). SFB-specific T cells activated in pregnant female mice following poly(I:C) injection can also contribute to IL-17A-mediated neurodevelopmental abnormalities in mouse offspring (Kim et al., 2017). Whether the same or different cytokine requirements apply to SFB-induced Th17 cells in the ileum at steady state and at distal sites during inflammation remains to be determined. It is still unclear whether humans harbor SFB, although a recent posted report provides evidence that some infants may have small amounts of a related species (Jonsson et al., 2019 in BioRxiv). It will be of considerable interest to determine if such bacteria in humans influence Th17 cell differentiation, activation, and trafficking.

## Acknowledgments

We thank Ken Cadwell, Sang V. Kim, Jason A. Hall, and Lin Wu for valuable discussion, and Biogen for providing control-Ig and LTβR-Ig. This work was supported by fellowships from the TOYOBO Bioscience foundation (T.S.), Human Frontier Science Program (T.S.), UIC start-up funds (T.S.), Schweppe Award in Translational Research (T.S.), the Helen and Martin Kimmel Center for Biology and Medicine (D.R.L.), The Judith and Stewart Colton Center for Autoimmunity (D.R.L), NYSTEM award (D.R.L), National Institutes of Health grant R01DK103358 (D.R.L.), and by the Howard Hughes Medical Institute (D.R.L.)

## Author Contributions

TS designed and performed most experiments and analyzed the data. TK performed core experiments and analyzed the data with TS. VF contributed on performing immunohistochemistry with JT and SRS. RK established *ex vivo* priming assay and collected data. YY and AC assisted with experiments. YY generated TCR transgenic mice. CN helped TS to maintain SFB-free animal colonies. SRS provided instructive suggestions about Th17 cell migration. TS and DRL wrote the manuscript. DRL supervised the research and participated in experimental design.

## Declaration of Interests

D.R.L consults and has equity interest in Chemocentryx, Vedanta Biosciences, and Pfizer Phamaceuticals.

## STAR Methods

### KEY RESOURCES TABLE

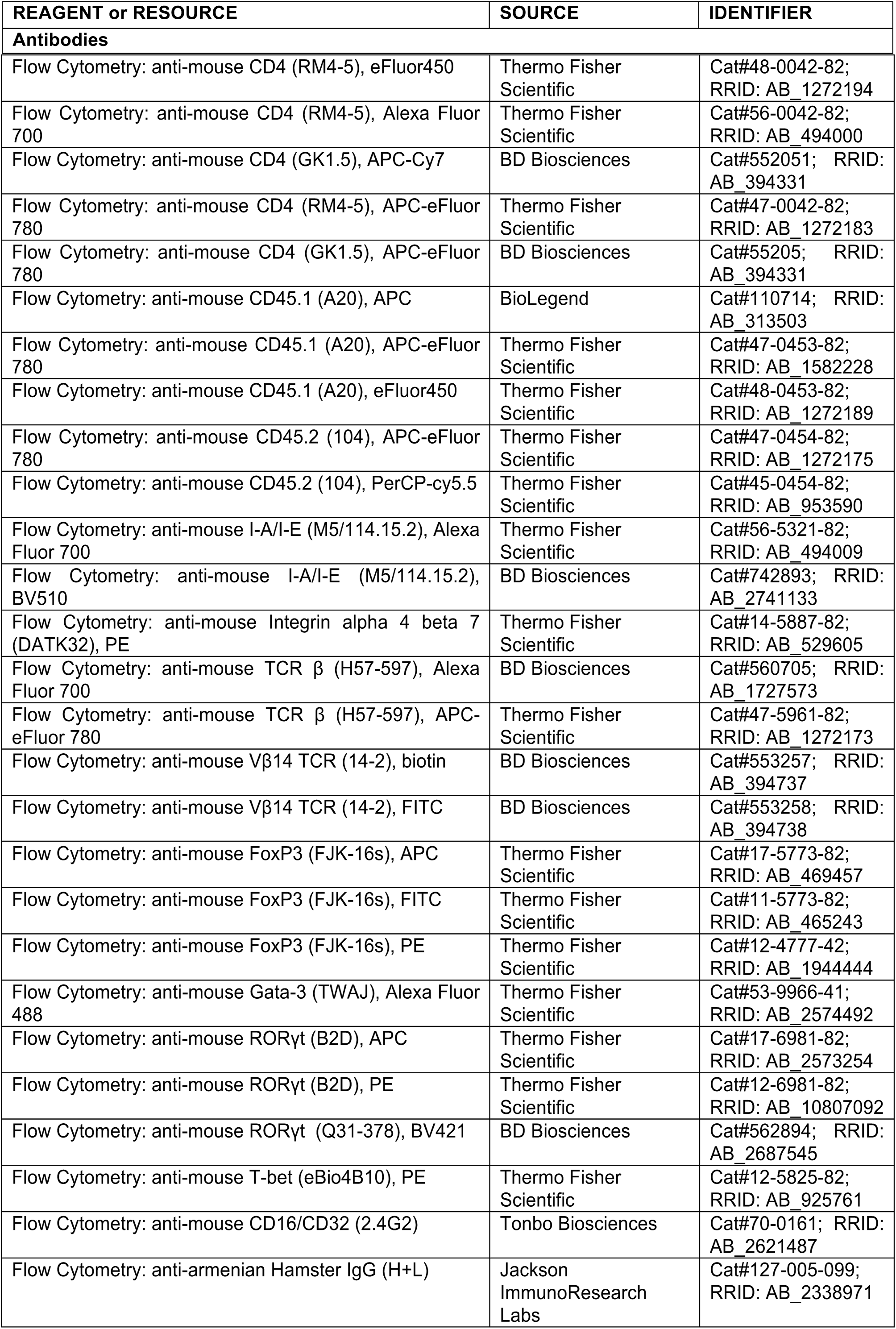
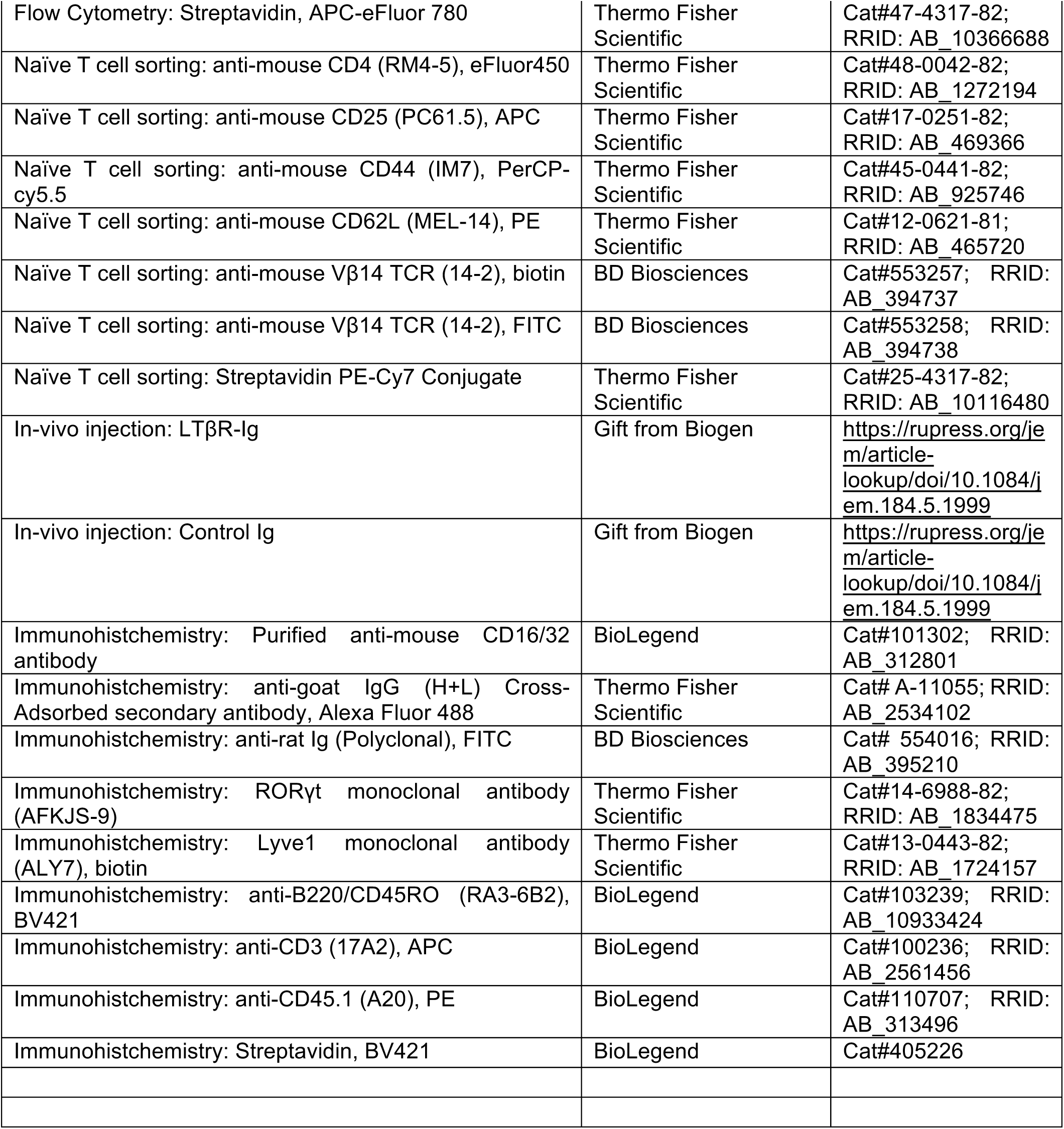

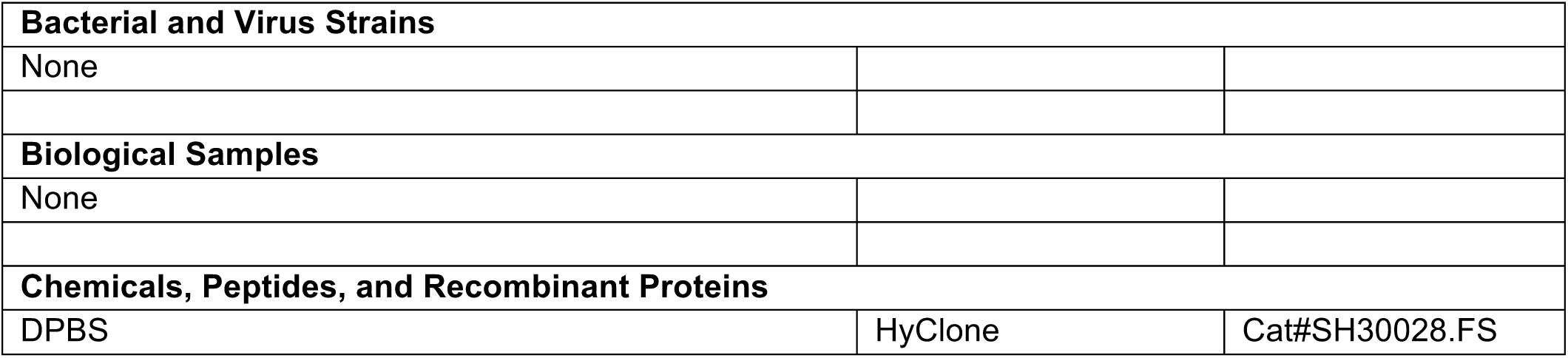
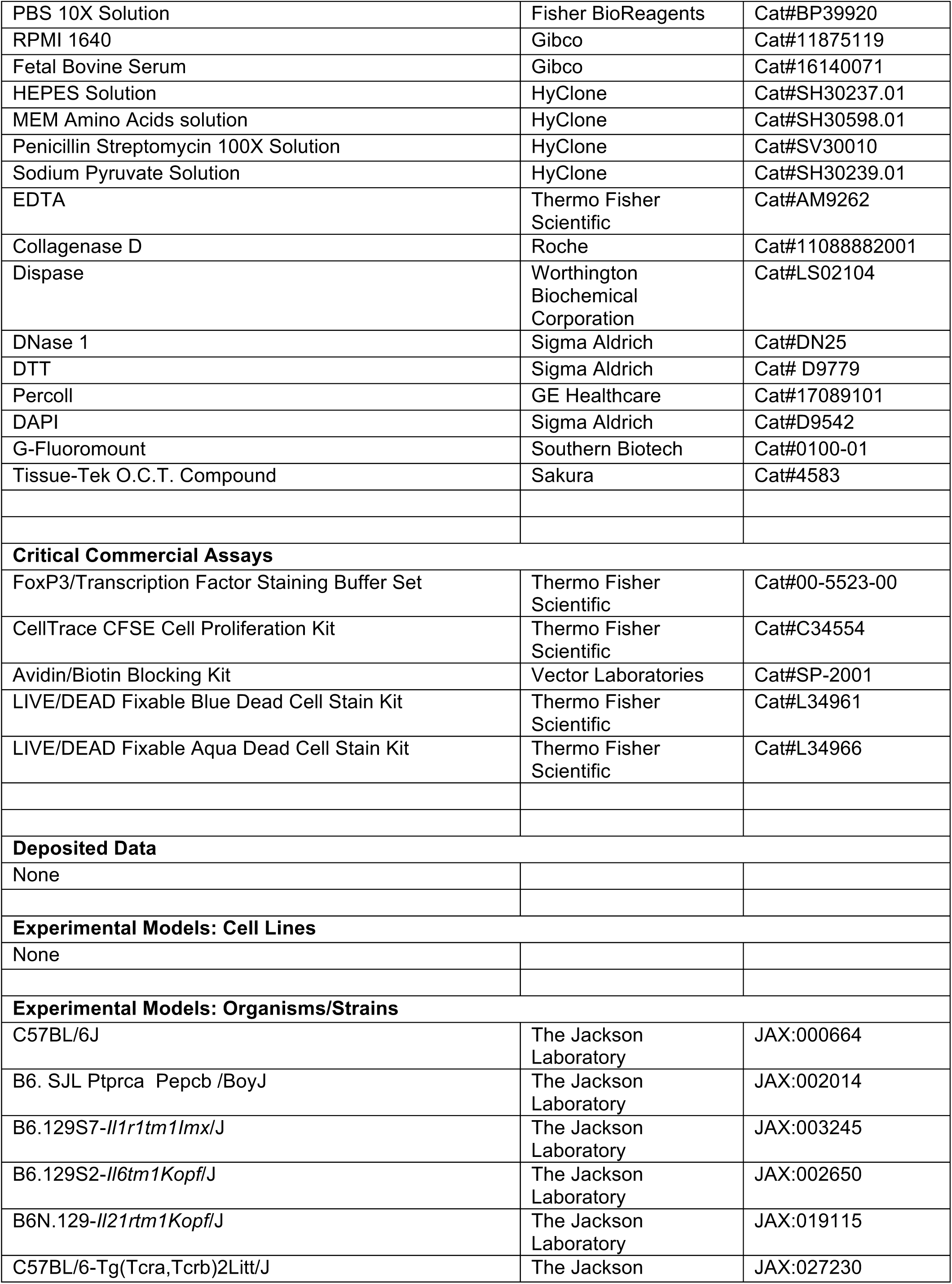
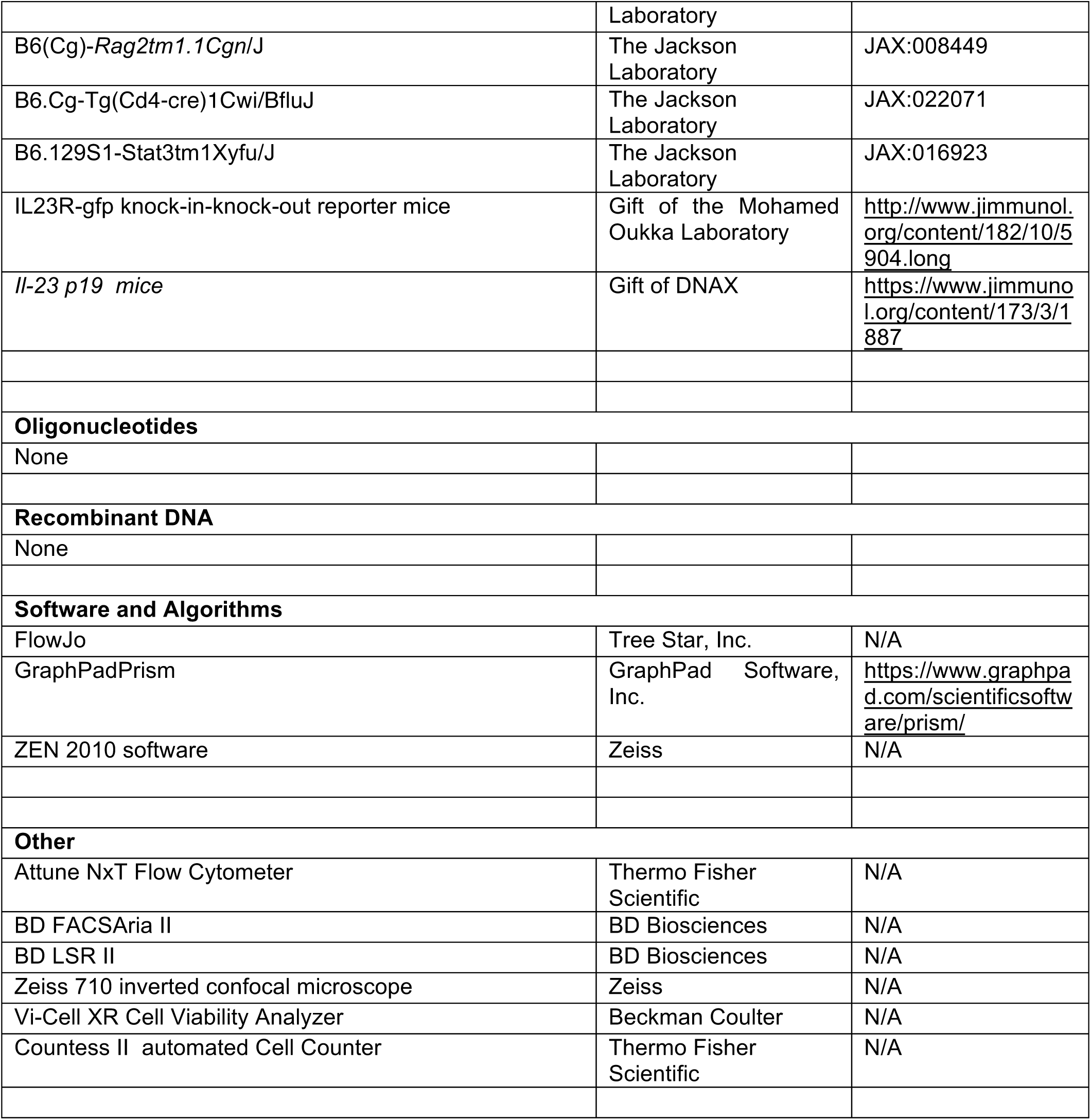

### Mouse Strains and Vivarium Housing

C57BL/6J mice were obtained from the Jackson Laboratory as SFB-negative mice. All transgenic animals were bred and maintained in the animal facility of the Skirball Institute (NYU School of Medicine) and Bio Research Laboratory of University of Illinois at Chicago (UIC) under specific-pathogen-free (SPF) conditions. To maintain SFB-free mice, autoclaved cages, autoclaved acidified water, and fresh chow were used and all cages were changed weekly by investigators. *Il-23r*^*gfp*^ mice (Awasthi et al., 2009) were provided by M. Oukka and maintained in *Rag2*^*-/-*^ (JAX) background. SFB-specific Th17-TCRTg (7B8) mice were previously described (Yang et al., 2014) and maintained on the Ly5.1 *(Cd45.1)* background (JAX; B6. SJL-Ptprca Pepcb/BoyJ). Itgb7 mice were purchased from JAX and maintained on the Ly5.1 *(Cd45.1)* or Ly5.2 *(Cd45.2)* background. *Il-23 p19*^*- /-*^ mice were obtained from DNAX (Lieberman et al., 2004) and maintained in SFB-free condition by treatment with ampicillin using autoclaved cages for 2 weeks. *Il-1r1* (JAX), *Il-6* (JAX), and *Il-21r* (JAX) mice were maintained with Jackson flora (SFB negative). Six-to 16-week old animals were used for *in vivo* experiments. All animal procedures were in accordance with protocols approved by the Institutional Animal Care and Use Committee of the NYU School of Medicine and UIC College of Medicine. All phenotypes were compared using littermate controls and the results were combined in most of the experiments.

### Acute SFB Colonization

Fecal pellets were collected from SFB-enriched *Il-23r Rag2* DKO mice (SFB), or SFB-free JAX B6 mice (Jax). Fresh fecal pellets were homogenized in ice-cold PBS, passed through a 100 μm filter, pelleted at 3400 rpm at 4ΰC for 10 minutes, and re-suspended in ice-cold PBS. Each SFB-free animal was administered 1/4 pellet by oral gavage. Before mice were used for the gavage experiments, we confirmed that they were SFB negative by qPCR, as described in (Sano et al., 2015). The state of SFB colonization in gavaged mice was confirmed by microscopic observation of the disrupted epithelial fraction during gut preparation or by qPCR. Littermates were cohoused from their birth or after SFB gavage in most of the experiments.

### SFB-specific Transgenic T Cell Transfer

SFB-enriched fecal extracts were administered by oral gavage to SFB-free mice 3 or 4 days before T cell transfer. 7B8 naïve T cells were purified as DAPI^-^, CD4^+^, CD62L^hi^, CD44^low^, CD25^-^, TCRVβ14^+^ T cells from spleen and lymph nodes (except MLN) of 7B8 TCR Tg mice using the Aria II cell sorter (BD). To monitor proliferation, sorted 7B8 naïve T cells were stained with CellTrace CFSE Cell Proliferation kit (Thermo Fisher Scientific). After cell numbers were calculated, 5,000 or 50,000 naïve 7B8 T cells were administered to recipient mice by intravenous retro-orbital injection. Animals were sacrificed and analyzed at multiple time points following naïve T cell transfer. Mononuclear cells from multiple tissues such as MLN, PP, spleen, duodenum, ileum, and colon were harvested as described (Yang et al., 2014).

### Depletion of PPs from Mouse Small Intestine

To abrogate development of gut-associated PPs, we intravenously injected SFB-free pregnant C57BL/6 mice with 150 μg of LTβR-Ig on gestational days 16.5 and 18.5, as described (Rennert et al., 1996). As an experimental negative control, control isotype-matched IgG was intravenously injected. During the gut prep, the number of PPs and MLNs were isolated and counted.

### Isolation of Lamina Propria Lymphocytes

Mesenteric fat tissue and Peyer’s patches were carefully removed from intestinal tissues. The proximal and distal 1/3 of small intestine were designated as duodenum and ileum, respectively. Tissues were incubated in 5 mM EDTA in PBS containing freshly prepared 1 mM of DTT for 20 min at 37°C with rotation (200-250 rpm). During the step, we confirmed that the treated tissues were not stacked at the bottom of the tubes. After this 1^st^ wash step, SFB attaching to epithelial cells can be detected under the microscopic observation in the supernatant (Epithelial rich fraction). Remaining tissues were incubated in a second EDTA wash (5 mM EDTA in PBS) for 10 min at 37°C with rotation (200-250 rpm). After tissues were washed with RPMI 10% FCS medium to remove EDTA, tissues were then further digested (RPMI media containing 10% fetal calf serum (FCS), 1.0 mg/ml each of Collagenase D (Roche) and 100 μg/ml DNase I (Sigma), and 0.01 U/ml Dispase (Worthington)) at 37°C for 30 min (duodenum and ileum) or 45 min (colon) with rotation (125-250 rpm). During the step, we confirmed that the treated tissues were not stacked at the bottom of the tubes. After the tissues were shaken vigorously in 50 ml tubes by hand for 30 seconds, 14 ml of ice cold RPMI 10% FCS was added to the tube. The digested tissues were then passed through a 70μm cell strainer. Mononuclear cells were enriched on 40:80 Percoll gradients by centrifugation (2200 rpm) for 22 min at room temperature. Lamina propria (LP) lymphocytes were collected from the Percoll gradient interphase. The recovered LP lymphocytes were washed with ice cold RPMI 10% FCS and the total cell numbers were counted by Vi-Cell XR Cell Viability Analyzer (Beckman Coulter) or Countess™ II automated Cell Counter (ThermoFisher).

### Cell Staining for Flow Cytometry

For analysis of T helper cell linages, live cells were stained with anti-CD3, anti-TCRβ, anti-CD4, and anti-MHC classII. To distinguish transferred 7B8 T cells from the host cells, live cells were stained with anti-TCRVβ14, anti-CD45.1, and anti-CD45.2. Live-dead labeling with DAPI (for live cell analysis) was performed and only live cells were analyzed. For intracellular protein staining, live-dead labeling with Aqua (Thermo Fisher Scientific) was performed. After extracellular proteins were labeled with antibodies, cells were fixed using fixation/permeabilization buffer (Thermo Fisher Scientific). Fixed cells were permeabilized using Permeabilization buffer (Thermo Fisher Scientific) and further stained with anti-Foxp3 and anti-RORγt using the Foxp3 staining buffer set (Thermo Fisher Scientific). Non-specific or Fc-receptor binding were blocked with control IgG and Fc-blocker during staining. Flow cytometric analysis was performed on a LSRII (BD Biosciences), a FACS Aria (BD Biosciences), or Attune (Thermo Fisher Scientific). All data were re-analyzed using FlowJo (Tree Star).

### Confocal microscopy

Mesenteric lymph nodes and Peyer’s patches were harvested, and fixed in 4% PFA for 1.5 hours at 22–25°C with gentle shaking. Organs were then dehydrated by sucrose gradient (15% sucrose in PBS for 1 hour at 4°C, then 30% sucrose at 4°C overnight), embedded in OCT (Sakura), and snap-frozen in dry ice-cooled 2-methylbutane. 8- to 10-μm sections were cut and air-dried at least 30 minutes. All staining was performed at 22– 25°C. Sections were permeabilized for 30 min with 0.5% Triton X-100 in PBS, then incubated for 10–30 min in block buffer consisting of 0.1% Triton X-100 in PBS with 1% BSA (mass/vol) and 10% normal donkey serum. An Avidin/Biotin blocking kit (Vector Laboratories) was used with biotinylated antibodies. Stains were done with antibodies diluted in the block buffer. The following fluorochrome-conjugated antibodies were used: anti-CD45.1 (A20), anti-CD3 (17A2) anti-B220/CD45RO (RA3-6B2), and streptavidin (all from BioLegend), Biotin-conjugated anti-Lyve1 (ALY7, 2.5 μg/ml) was from Thermo Fisher Scientific.

For RORγt detection, primary anti-RORγt antibodies (1:30, clone AFKJS-9, Thermo Fisher Scientific 14-6988) were detected with secondary FITC Goat Anti-Rat Ig (1:500, BD Biosciences 554016), then tertiary Alexa Fluor® 488 donkey anti-goat (1:500, Thermo Fisher Scientific A11055). 10 μg/ml anti-CD16/32 (BioLegend, clone 93) was used in the buffer for tertiary staining. Slides were mounted with G-Fluoromount (Southern Biotech) and visualized using a Zeiss 710 inverted confocal microscope with a 25× or 40× oil-immersion objective and ZEN 2010 software. For all direct comparisons samples were stained on the same day and imaged with the same settings.

### *Ex-vivo* priming assay

*Cd45.1/Cd45.1* WT B6 mice (8 week old females) were gavaged with SFB for two sequential days and MLN (L1: closest from cecum: highly associated to ileum) and PPs were isolated at several time points following the first gavage. *Cd45.2/Cd45.2* WT B6 mice were also prepared as SFB-negative control and their MLN and PPs were collected in the same way. Collected tissues from SFB negative and positive mice were combined and digested together with RPMI 10% FCS containing 100 μg/ml DNase I (Sigma), 1.0 mg/ml of Collagenase D (Roche) and 0.01 U/ml of Dispase (Worthington)) at 37°C for 30 min by gently shaking. Digested tissues were passed through 100 μm filter then stained for cell sorting. APCs were sorted following the gating. APCs from SFB gavaged: DAPI-, Dump^-^ (CD45R^-^, TCRβ^-^, NK1.1^-^ Ly6G^-^), CD45.1^+^, and CD11b^+^ or CD11c^+^ singlet cells, APCs from SFB negative mice: Dump^-^ (CD45R^-^, TCRβ^-^, NK1.1^-^ Ly6G^-^), CD45.2^+^, and CD11b^+^ or CD11c^+^ singlet cells. APCs were sorted from indicated whole tissues and incubated with naïve 7B8 T cells with IMDM 10% FCS for 24 h. T cell activation was assessed by flow cytometry after staining for TCRvβ14, CD4, CD25, CD69, and CD62L.

### Statistics

Two-tailed unpaired Student’s t-tests were performed to compare the results using Excel and Prism. We treated less than 0.05 of P-value as significant differences. * p<0.05, ** p<0.01, *** p<0.001, and **** p<0.0001. N.S.: No Significant.

## Supplemental Information

### Supplemental Figure Legends

**Figure S1.**
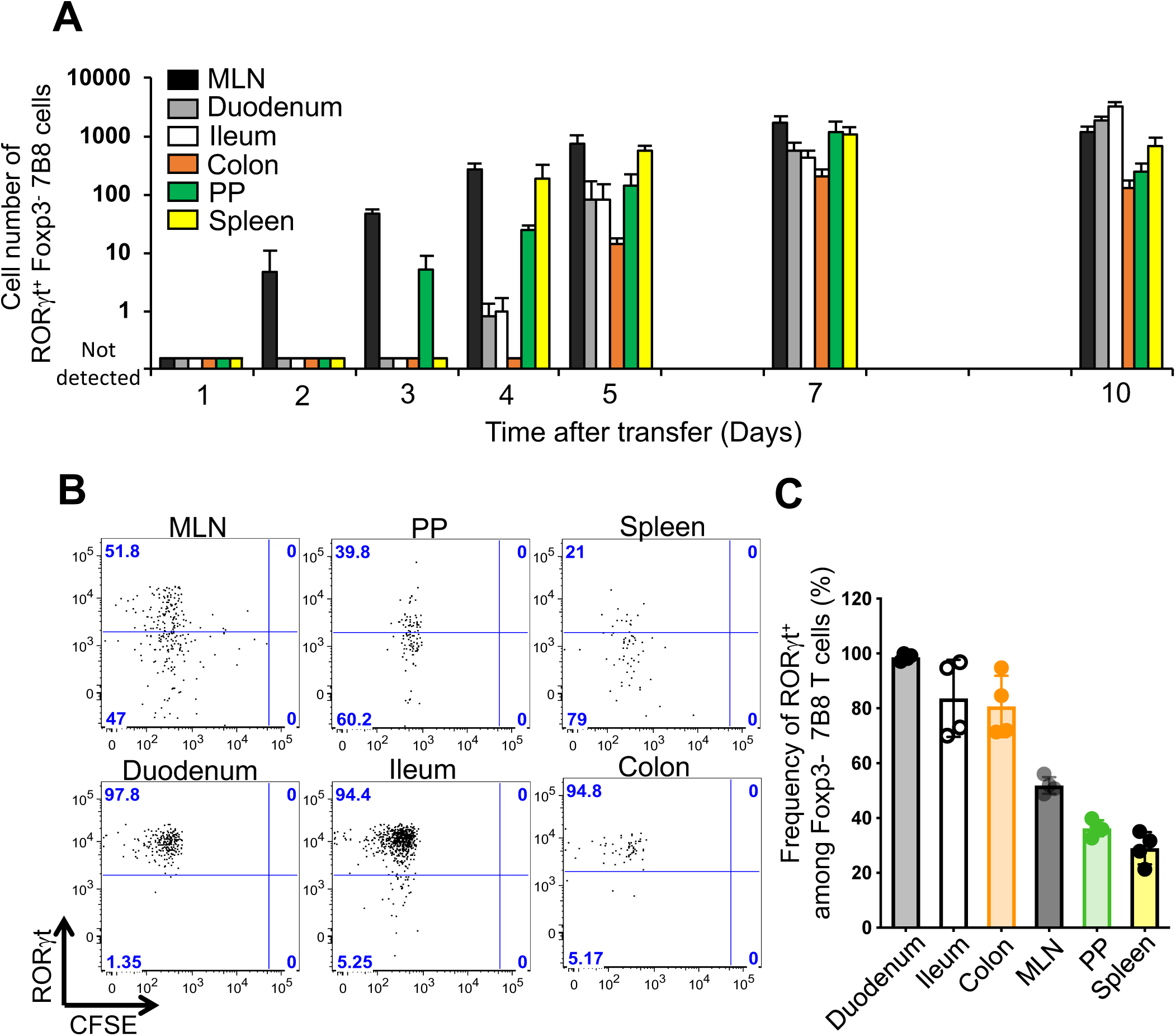
Kinetics of SFB-specific Th17 cell differentiation in tissues of SFB-colonized mice (related to Figure 1) **(A)** Kinetics of SFB-specific Th17 cell differentiation following transfer of 5,000 naïve 7B8 T cells (*Cd45.1/ Cd45.2)*. The number of SFB-specific Th17 cells in the MLN, duodenum, ileum, colon, PPs, and spleens of SFB-gavaged B6 mice (*Cd45.2/ Cd45.2)* is shown. SFB-specific Th17 cells were defined as Foxp3^-^ RORγt^+^ 7B8 T cells. All data were generated from one time course experiment with 3 or 4 mice at each time point, but multiple time points were repeated independently with similar results. Data are represented as mean +/- SD. **(B)** CFSE dilution and RORγt expression among the transferred 7B8 T cells in MLN, duodenum, ileum, colon, PPs, and spleens of SFB-gavaged mice at 10 days following transfer, representative of data in Fig. S1A. **(C)** Frequency of RORγt^+^ cells among Foxp3^-^ 7B8 T cells in the described location. Each symbol represents a single mouse, and the color bar indicates average +/- SD.

**Figure S2.**
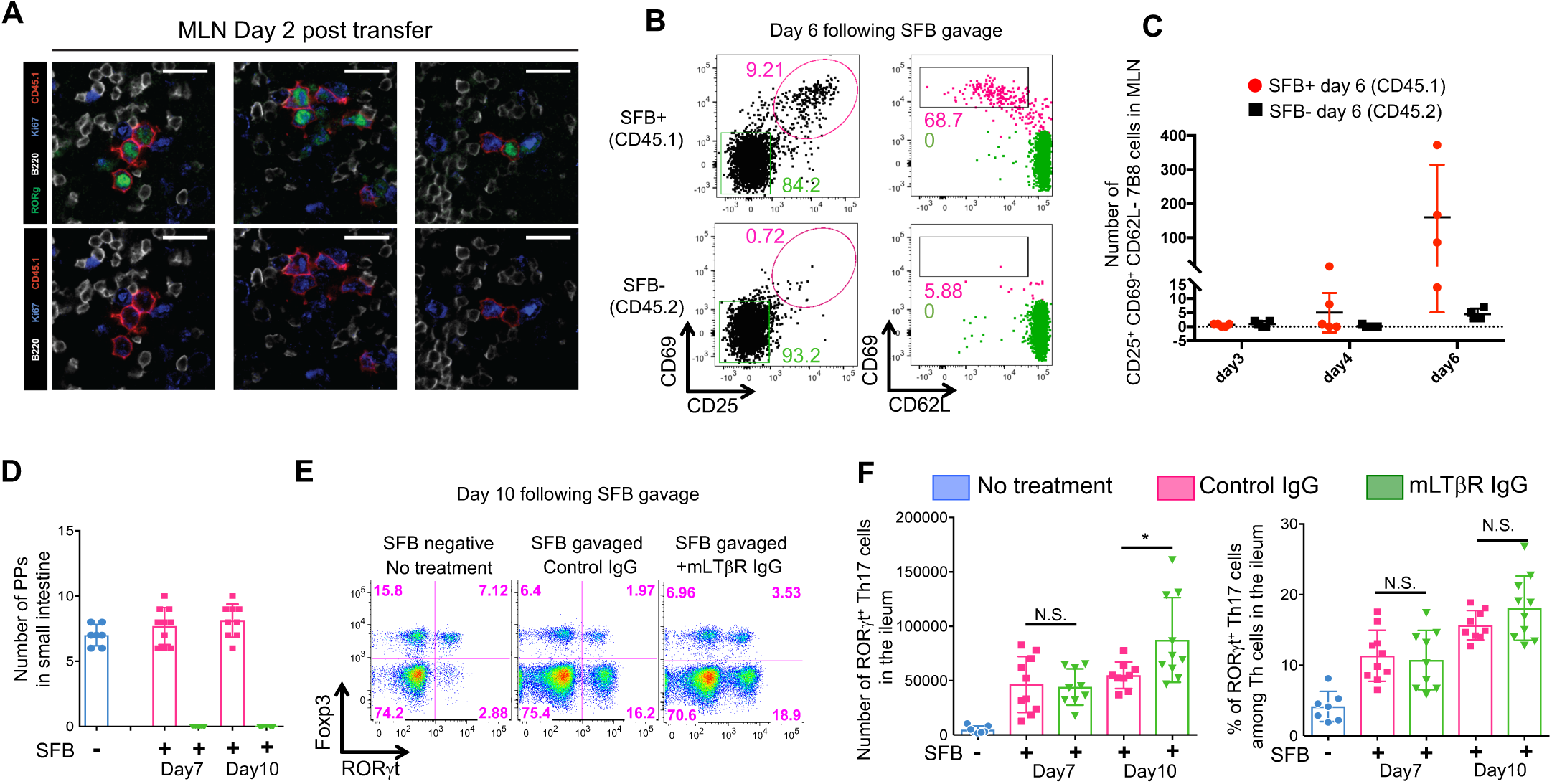
Role of MLN and PPs in Th17 cell induction following SFB gavage (related to Figure 2) **(A)** Proliferation of SFB-specific T cells in cell clusters within the MLN T cell zone and follicular border at 2 days following transfer of 50,000 naïve 7B8 T cells. Sections were stained with anti-RORγt^+^, anti-CD45.1, anti-B220, and anti-Ki67, and were analyzed by confocal microscopy. Representative of 5 mice from 2 experiments. **(B and C)** Ex vivo priming of naïve SFB-specific T cells by purified APCs (CD11b^+^ and CD11c^+^ cells). APCs from MLN were purified at several time points (Day3: n=5, Day4: n=5, Day6: n=4) after SFB or saline gavage and were co-cultured with 10,000 sorted naïve T cells from SFB-specific TCR Tg mice for 24 h. Activation was assessed by expression of CD25 and CD69 on cells gated for TCRvβ14 and CD4 expression (B). (C) Kinetics of antigen presentation activity by APCs from MLNs following SFB gavage. Activated SFB-specific T cells were defined as TCRvβ14^+^, CD4^+^, CD25^+^, CD69^+^, and CD62L^-^. **(D-F)** Th17 cell differentiation in PP-depleted mice gavaged with SFB. 150 μg LTβR-Ig were intravenously injected SFB-free pregnant B6 mice on gestational days 16.5 and 18.5 (Rennert et al., 1996). The number of PPs in the small intestine after LTβR-IgG or control IgG administration during gestation (D). The frequency and total cell number of RORγt^+^ Foxp3^-^ Th17 cells in the ileum at day 10 after SFB gavage, with representative FACS panels (E) and cumulative data (F). Each symbol represents a single mouse, and the color bar indicates average +/- SD. Results from two independent experiments were combined (SFB negative: n=7, control IgG injected SFB-gavaged mice: n=10 (Day7) and 9 (Day10), LTβR-Ig injected SFB gavaged mice n=9 (Day7) and 10 (Day10). * p<0.05. N.S.: Not significant.

**Figure S3.**
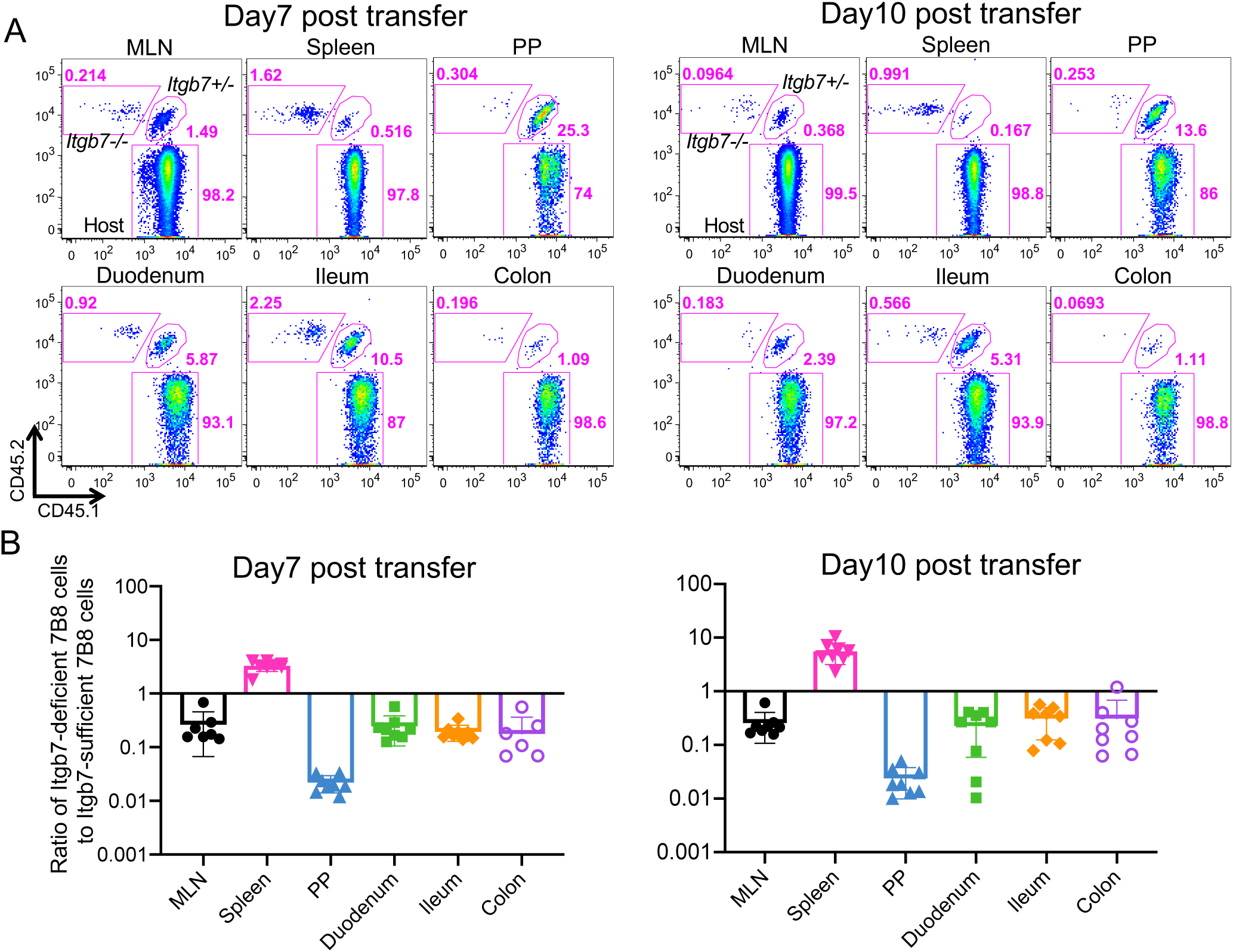
The role of α4β7 integrin in migration of SFB-specific Th17 cells from MLN to local tissues (related to Figure 3) **(A and B)** SFB-specific T cells in different tissues at 7 and 10 days following co-transfer of 5,000 naïve integrin β7 (*Itgb7*)-sufficient (*Cd45.1/Cd45.2*) and 5,000 *Itgb7*-deficient (*Cd45.2/Cd45.2*) 7B8 T cells into SFB gavaged mice (*Cd45.1/Cd45.1*). In (B), each symbol represents tissue cell ratio in a single mouse, and the color bar indicates average +/- SD. Two experiments were performed and results were combined (n=8 for each time point).

**Figure S4.**
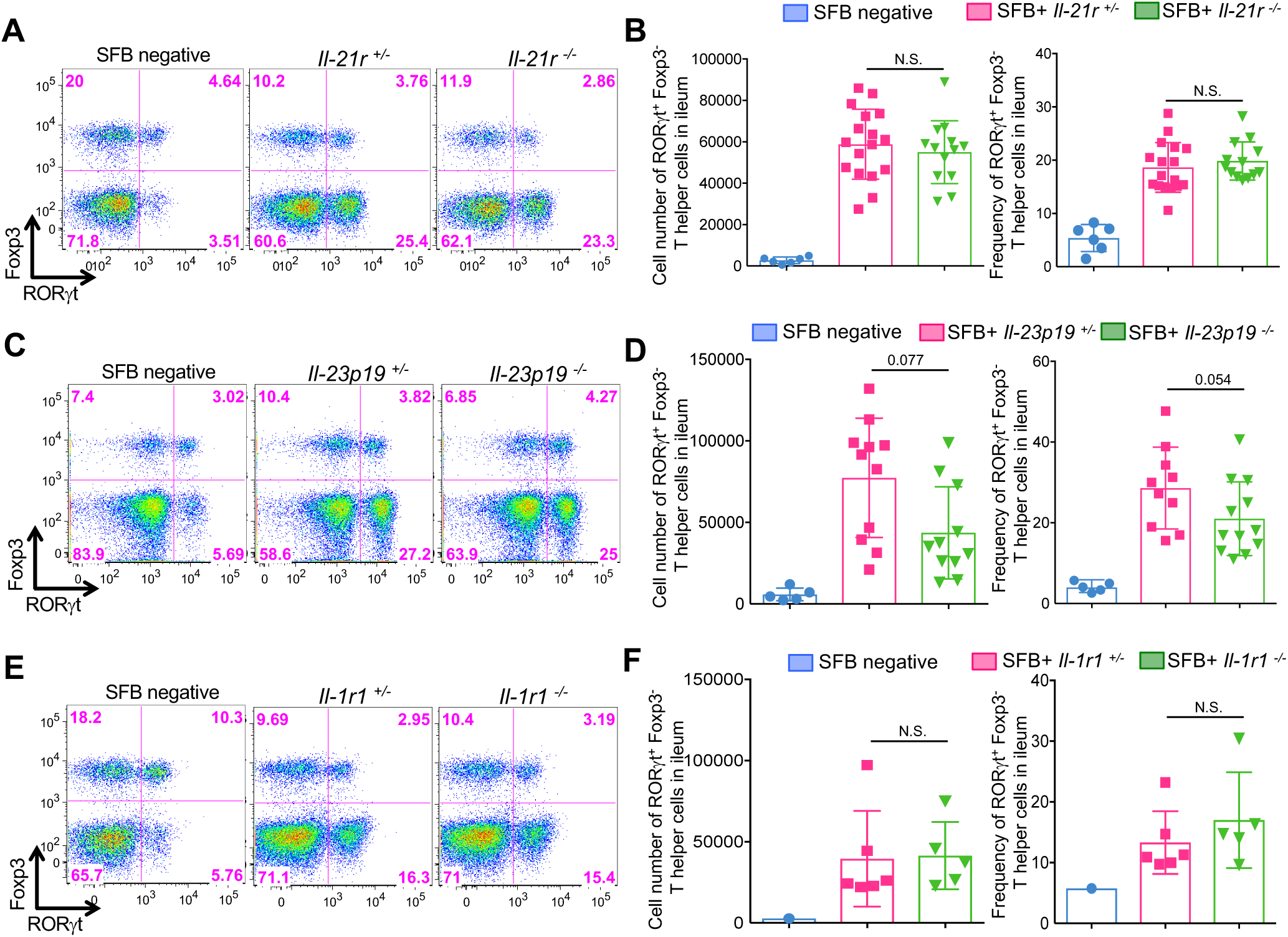
Cytokine requirements for SFB-induced Th17 cell differentiation *in vivo* (related to Figure 4) **(A-F)** Th17 cell differentiation in response to SFB gavage in mice deficient for IL-21R (A and B), IL-23 p19 (C and D), and IL1R1 (E and F). SFB was gavaged into *Il-21r, Il-23 p19*, and *Il1r1* sufficient and deficient co-housed littermates and at days 7 (IL-21R and IL1R1) or 10 (IL-23 p19) ileal LP RORγt^+^ Foxp3^-^ Th17 cells were quantified by FACS. (A), (C), and (E) are representative FACS panels; (B), (D), and (F) show aggregate Th17 cell numbers. Experiments were done at least twice with several mutant and littermate controls. Results were combined and each dot shows an individual mouse, with color bars indicating average +/- SD. N.S.: not significant.

**Figure S5.**
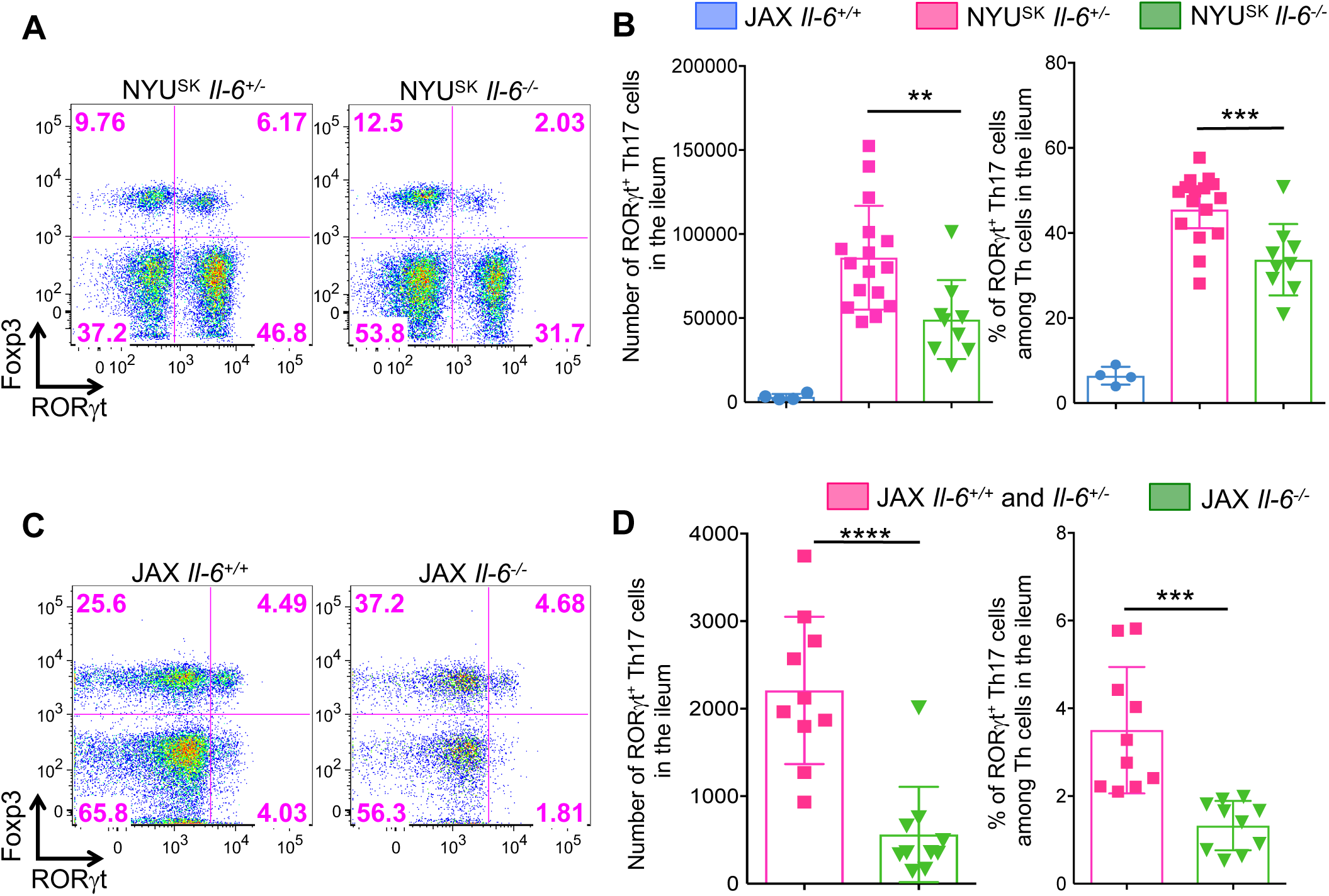
Comparison of IL-6 requirements for Th17 cell differentiation in mice housed in vivaria with and without SFB (Related to Figure 4) **(A and B)** Th17 cells in ileum of stably SFB-colonized *Il-6* heterozygous or homozygous null C57BL/6 mice housed in the Skirball animal facility at New York University (NYU^SK^). Mice were confirmed as SFB^+^ by qPCR. Representative flow cytometry analysis (A) and statistics of number (B, left) and frequency (B, right) of RORγt^+^ Foxp3^-^ Th17 cells in the ileum. Experiments were done three times with biological replicates (SFB negative *Il-6*^*+/-*^: n=4, *Il-6*^*+/-*^: n=16, and *Il-6*^*-/-*^: n=9) and the results were combined. Littermates were cohoused from birth. **(C and D)** IL-6 requirement for Th17 cell differentiation in mice housed at Jackson Laboratory (JAX) in SFB-free conditions. Data are shown as in (A and B). Experiments were done twice with 10 mice total in each group, and combined results are shown. Dots show individual mice and color bars indicate average +/- SD. ** p<0.01, *** p<0.001, and **** p<0.0001. N.S.: not significant.

**Figure S6.**
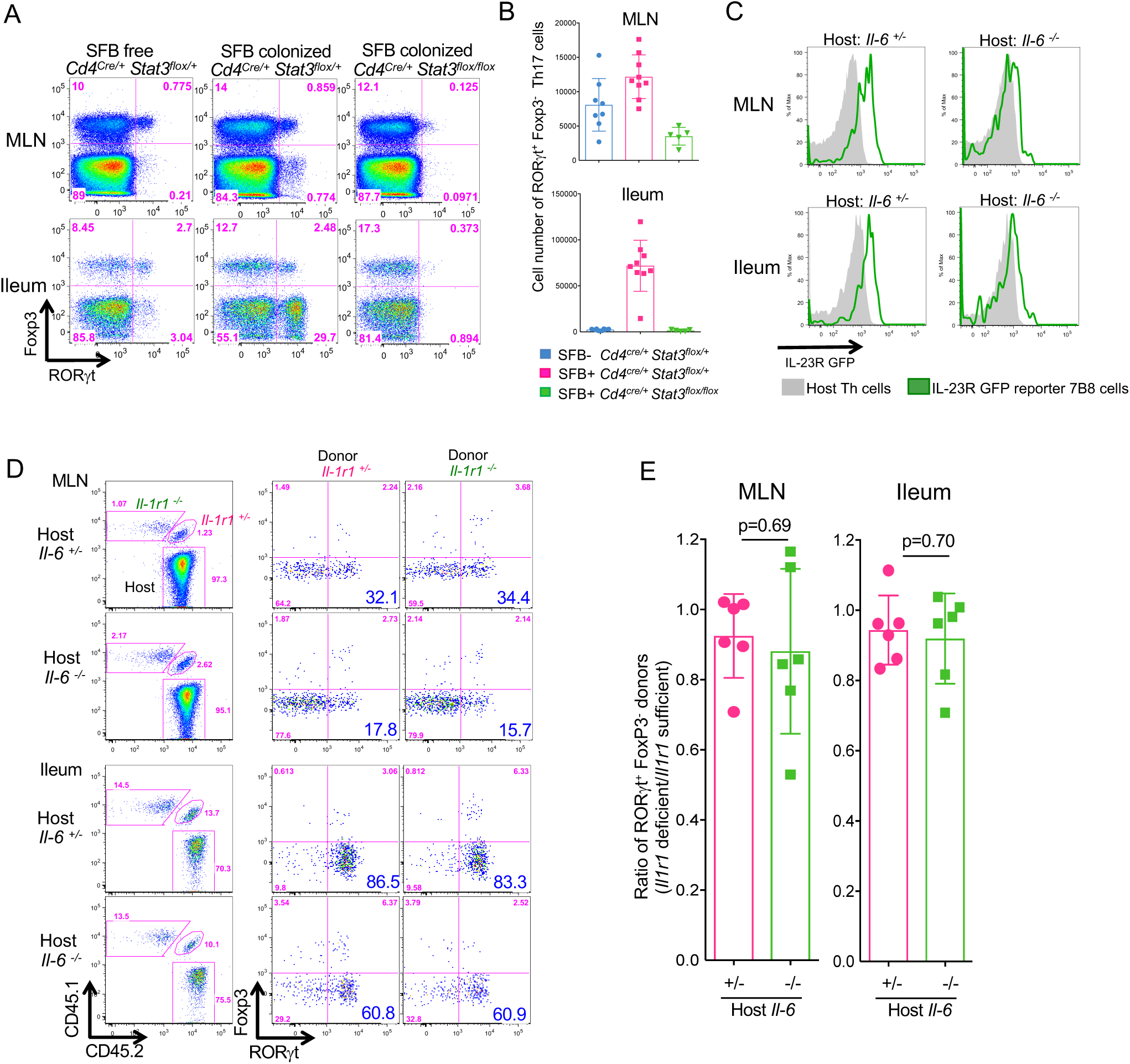
Stat3-dependent cytokine redundancy in SFB-specific Th17 cell differentiation in the MLN and ileum (related to figure 5) **(A-B)** Th17 cells from the MLN and ileum of *Cd4*^*cre/+*^ *Stat3*^*flox/+*^ and *Cd4*^*cre/+*^ *Stat3*^*flox/flox*^ mice stably colonized with SFB or SFB-free. Representative FACS plots (A) and numbers of RORγt^+^Foxp3^-^ CD4^+^ T cells (B). Experiments were repeated three times with several mice and combined data are shown. Dots represent individual mice and the color bars indicate average +/- SD. (SFB negative control: n=9, *Cd4*^*cre/+*^ *Stat3*^*flox/+*^: n=9, *Cd4*^*cre/+*^ *Stat3*^*flox/flox*^: n=5). **(C)** Expression of *Il-23r*^*gfp*^ reporter in SFB-specific 7B8 TCR transgenic Th17 cells in the MLN and the ileum of *Il-6* sufficient and deficient mice at 7 days following naïve T cell transfer. **(D and E)** SFB-specific Th17 cell differentiation in the presence or absence of IL-1R1 signaling in *Il-6* sufficient and deficient mice. 5000 naïve T cells from cytokine receptor sufficient and deficient 7B8 Tg mice, with indicated isotypes, were co-transferred into SFB-gavaged *Il-6* sufficient and deficient mice (*Cd45.2/Cd45.2*). Representative FACS plots of ileal T cells at 7 days following naïve T cell transfer (D). Ratios of RORγt^+^ Foxp3^-^ Th17 cells (*Il1r1* sufficient vs deficient T cells) in *Il-6* sufficient (n=6) and deficient (n=6) mice (E). Data from two experiments with similar results were combined. Dots represent individual mice and color bars indicates average +/- SD.

## Notes

### Competing Interest Statement

The authors have declared no competing interest.

## References

Acosta-Rodriguez, E.V., Napolitani, G., Lanzavecchia, A., and Sallusto, F. (2007). Interleukins 1beta and 6 but not transforming growth factor-beta are essential for the differentiation of interleukin 17-producing human T helper cells. Nat Immunol 8, 942–949.

Agnello, D., Lankford, C.S., Bream, J., Morinobu, A., Gadina, M., O’Shea, J.J., and Frucht, D.M. (2003). Cytokines and transcription factors that regulate T helper cell differentiation: new players and new insights. J Clin Immunol 23, 147–161.

Atarashi, K., Nishimura, J., Shima, T., Umesaki, Y., Yamamoto, M., Onoue, M., Yagita, H., Ishii, N., Evans, R., Honda, K., and Takeda, K. (2008). ATP drives lamina propria T(H)17 cell differentiation. Nature 455, 808–812.

Awasthi, A., Riol-Blanco, L., Jager, A., Korn, T., Pot, C., Galileos, G., Bettelli, E., Kuchroo, V.K., and Oukka, M. (2009). Cutting edge: IL-23 receptor gfp reporter mice reveal distinct populations of IL-17-producing cells. J Immunol 182, 5904–5908.

Bettelli, E., Carrier, Y., Gao, W., Korn, T., Strom, T.B., Oukka, M., Weiner, H.L., and Kuchroo, V.K. (2006). Reciprocal developmental pathways for the generation of pathogenic effector TH17 and regulatory T cells. Nature 441, 235–238.

Bradley, C.P., Teng, F., Felix, K.M., Sano, T., Naskar, D., Block, K.E., Huang, H., Knox, K.S., Littman, D.R., and Wu, H.J. (2017). Segmented Filamentous Bacteria Provoke Lung Autoimmunity by Inducing Gut-Lung Axis Th17 Cells Expressing Dual TCRs. Cell Host Microbe 22, 697–704 e694.

Chen, Y., Langrish, C.L., McKenzie, B., Joyce-Shaikh, B., Stumhofer, J.S., McClanahan, T., Blumenschein, W., Churakovsa, T., Low, J., Presta, L., et al. (2006). Anti-IL-23 therapy inhibits multiple inflammatory pathways and ameliorates autoimmune encephalomyelitis. J Clin Invest 116, 1317–1326.

Chudnovskiy, A., Mortha, A., Kana, V., Kennard, A., Ramirez, J.D., Rahman, A., Remark, R., Mogno, I., Ng, R., Gnjatic, S., et al. (2016). Host-Protozoan Interactions Protect from Mucosal Infections through Activation of the Inflammasome. Cell 167, 444–456 e414.

Chung, Y., Chang, S.H., Martinez, G.J., Yang, X.O., Nurieva, R., Kang, H.S., Ma, L., Watowich, S.S., Jetten, A.M., Tian, Q., and Dong, C. (2009). Critical regulation of early Th17 cell differentiation by interleukin-1 signaling. Immunity 30, 576–587.

Cua, D.J., Sherlock, J., Chen, Y., Murphy, C.A., Joyce, B., Seymour, B., Lucian, L., To, W., Kwan, S., Churakova, T., et al. (2003). Interleukin-23 rather than interleukin-12 is the critical cytokine for autoimmune inflammation of the brain. Nature 421, 744–748.

Erle, D.J., Briskin, M.J., Butcher, E.C., Garcia-Pardo, A., Lazarovits, A.I., and Tidswell, M. (1994). Expression and function of the MAdCAM-1 receptor, integrin alpha 4 beta 7, on human leukocytes. J Immunol 153, 517–528.

Gaffen, S.L., Jain, R., Garg, A.V., and Cua, D.J. (2014). The IL-23-IL-17 immune axis: from mechanisms to therapeutic testing. Nat Rev Immunol 14, 585–600.

Geem, D., Medina-Contreras, O., McBride, M., Newberry, R.D., Koni, P.A., and Denning, T.L. (2014). Specific microbiota-induced intestinal Th17 differentiation requires MHC class II but not GALT and mesenteric lymph nodes. J Immunol 193, 431–438.

Ghoreschi, K., Laurence, A., Yang, X.P., Tato, C.M., McGeachy, M.J., Konkel, J.E., Ramos, H.L., Wei, L., Davidson, T.S., Bouladoux, N., et al. (2010). Generation of pathogenic T(H)17 cells in the absence of TGF-beta signalling. Nature 467, 967–971.

Goto, Y., Panea, C., Nakato, G., Cebula, A., Lee, C., Diez, M.G., Laufer, T.M., Ignatowicz, L., and Ivanov, II (2014). Segmented filamentous bacteria antigens presented by intestinal dendritic cells drive mucosal Th17 cell differentiation. Immunity 40, 594–607.

Hall, J.A., Bouladoux, N., Sun, C.M., Wohlfert, E.A., Blank, R.B., Zhu, Q., Grigg, M.E., Berzofsky, J.A., and Belkaid, Y. (2008). Commensal DNA limits regulatory T cell conversion and is a natural adjuvant of intestinal immune responses. Immunity 29, 637–649.

Harris, T.J., Grosso, J.F., Yen, H.R., Xin, H., Kortylewski, M., Albesiano, E., Hipkiss, E.L., Getnet, D., Goldberg, M.V., Maris, C.H., et al. (2007). Cutting edge: An in vivo requirement for STAT3 signaling in TH17 development and TH17-dependent autoimmunity. J Immunol 179, 4313–4317.

Honda, K., and Littman, D.R. (2016). The microbiota in adaptive immune homeostasis and disease. Nature 535, 75–84.

Hooper, L.V., Littman, D.R., and Macpherson, A.J. (2012). Interactions between the microbiota and the immune system. Science 336, 1268–1273.

Ivanov, II, Atarashi, K., Manel, N., Brodie, E.L., Shima, T., Karaoz, U., Wei, D., Goldfarb, K.C., Santee, C.A., Lynch, S.V., et al. (2009). Induction of intestinal Th17 cells by segmented filamentous bacteria. Cell 139, 485–498.

Ivanov, II, Frutos Rde, L., Manel, N., Yoshinaga, K., Rifkin, D.B., Sartor, R.B., Finlay, B.B., and Littman, D.R. (2008). Specific microbiota direct the differentiation of IL-17-producing T-helper cells in the mucosa of the small intestine. Cell Host Microbe 4, 337–349.

Ivanov, II, McKenzie, B.S., Zhou, L., Tadokoro, C.E., Lepelley, A., Lafaille, J.J., Cua, D.J., and Littman, D.R. (2006). The orphan nuclear receptor RORgammat directs the differentiation program of proinflammatory IL-17+ T helper cells. Cell 126, 1121–1133.

Jain, A., Song, R., Wakeland, E.K., and Pasare, C. (2018). T cell-intrinsic IL-1R signaling licenses effector cytokine production by memory CD4 T cells. Nat Commun 9, 3185.

Johansson-Lindbom, B., and Agace, W.W. (2007). Generation of gut-homing T cells and their localization to the small intestinal mucosa. Immunol Rev 215, 226–242.

Jones, S.A., Scheller, J., and Rose-John, S. (2011). Therapeutic strategies for the clinical blockade of IL-6/gp130 signaling. J Clin Invest 121, 3375–3383.

Kamimura, D., Ishihara, K., and Hirano, T. (2003). IL-6 signal transduction and its physiological roles: the signal orchestration model. Rev Physiol Biochem Pharmacol 149, 1–38.

Kim, S., Kim, H., Yim, Y.S., Ha, S., Atarashi, K., Tan, T.G., Longman, R.S., Honda, K., Littman, D.R., Choi, G.B., and Huh, J.R. (2017). Maternal gut bacteria promote neurodevelopmental abnormalities in mouse offspring. Nature 549, 528–532.

Koopman, J.P., Stadhouders, A.M., Kennis, H.M., and De Boer, H. (1987). The attachment of filamentous segmented micro-organisms to the distal ileum wall of the mouse: a scanning and transmission electron microscopy study. Lab Anim 21, 48–52.

Korn, T., Bettelli, E., Gao, W., Awasthi, A., Jager, A., Strom, T.B., Oukka, M., and Kuchroo, V.K. (2007). IL-21 initiates an alternative pathway to induce proinflammatory T(H)17 cells. Nature 448, 484–487.

Korn, T., Bettelli, E., Oukka, M., and Kuchroo, V.K. (2009). IL-17 and Th17 Cells. Annu Rev Immunol 27, 485–517.

Langrish, C.L., Chen, Y., Blumenschein, W.M., Mattson, J., Basham, B., Sedgwick, J.D., McClanahan, T., Kastelein, R.A., and Cua, D.J. (2005). IL-23 drives a pathogenic T cell population that induces autoimmune inflammation. J Exp Med 201, 233–240.

Lecuyer, E., Rakotobe, S., Lengline-Garnier, H., Lebreton, C., Picard, M., Juste, C., Fritzen, R., Eberl, G., McCoy, K.D., Macpherson, A.J., et al. (2014). Segmented filamentous bacterium uses secondary and tertiary lymphoid tissues to induce gut IgA and specific T helper 17 cell responses. Immunity 40, 608–620.

Lee, J.Y., Hall, J.A., Kroehling, L., Wu, L., Najar, T., Nguyen, H.H., Lin, W.Y., Yeung, S.T., Silva, H.M., Li, D., et al. (2020). Serum Amyloid A Proteins Induce Pathogenic Th17 Cells and Promote Inflammatory Disease. Cell 180, 79–91 e16.

Lieberman, L.A., Cardillo, F., Owyang, A.M., Rennick, D.M., Cua, D.J., Kastelein, R.A., and Hunter, C.A. (2004). IL-23 provides a limited mechanism of resistance to acute toxoplasmosis in the absence of IL-12. J Immunol 173, 1887–1893.

Locci, M., Havenar-Daughton, C., Landais, E., Wu, J., Kroenke, M.A., Arlehamn, C.L., Su, L.F., Cubas, R., Davis, M.M., Sette, A., et al. (2013). Human circulating PD-1+CXCR3-CXCR5+ memory Tfh cells are highly functional and correlate with broadly neutralizing HIV antibody responses. Immunity 39, 758–769.

Maddur, M.S., Miossec, P., Kaveri, S.V., and Bayry, J. (2012). Th17 cells: biology, pathogenesis of autoimmune and inflammatory diseases, and therapeutic strategies. Am J Pathol 181, 8–18.

Manel, N., Unutmaz, D., and Littman, D.R. (2008). The differentiation of human T(H)-17 cells requires transforming growth factor-beta and induction of the nuclear receptor RORgammat. Nat Immunol 9, 641–649.

Mangan, P.R., Harrington, L.E., O’Quinn, D.B., Helms, W.S., Bullard, D.C., Elson, C.O., Hatton, R.D., Wahl, S.M., Schoeb, T.R., and Weaver, C.T. (2006). Transforming growth factor-beta induces development of the T(H)17 lineage. Nature 441, 231–234.

Mora, J.R., Bono, M.R., Manjunath, N., Weninger, W., Cavanagh, L.L., Rosemblatt, M., and Von Andrian, U.H. (2003). Selective imprinting of gut-homing T cells by Peyer’s patch dendritic cells. Nature 424, 88–93.

Mori, T., Miyamoto, T., Yoshida, H., Asakawa, M., Kawasumi, M., Kobayashi, T., Morioka, H., Chiba, K., Toyama, Y., and Yoshimura, A. (2011). IL-1beta and TNFalpha-initiated IL-6-STAT3 pathway is critical in mediating inflammatory cytokines and RANKL expression in inflammatory arthritis. Int Immunol 23, 701–712.

Nurieva, R., Yang, X.O., Martinez, G., Zhang, Y., Panopoulos, A.D., Ma, L., Schluns, K., Tian, Q., Watowich, S.S., Jetten, A.M., and Dong, C. (2007). Essential autocrine regulation by IL-21 in the generation of inflammatory T cells. Nature 448, 480–483.

Ohnmacht, C., Park, J.H., Cording, S., Wing, J.B., Atarashi, K., Obata, Y., Gaboriau-Routhiau, V., Marques, R., Dulauroy, S., Fedoseeva, M., et al. (2015). MUCOSAL IMMUNOLOGY. The microbiota regulates type 2 immunity through RORgammat(+) T cells. Science 349, 989–993.

Perrier, C., and Rutgeerts, P. (2011). Cytokine blockade in inflammatory bowel diseases. Immunotherapy 3, 1341–1352.

Rennert, P.D., Browning, J.L., Mebius, R., Mackay, F., and Hochman, P.S. (1996). Surface lymphotoxin alpha/beta complex is required for the development of peripheral lymphoid organs. J Exp Med 184, 1999–2006.

Sanchez-Munoz, F., Dominguez-Lopez, A., and Yamamoto-Furusho, J.K. (2008). Role of cytokines in inflammatory bowel disease. World J Gastroenterol 14, 4280–4288.

Sano, T., Huang, W., Hall, J.A., Yang, Y., Chen, A., Gavzy, S.J., Lee, J.Y., Ziel, J.W., Miraldi, E.R., Domingos, A.I., et al. (2015). An IL-23R/IL-22 Circuit Regulates Epithelial Serum Amyloid A to Promote Local Effector Th17 Responses. Cell 163, 381–393.

Schweighoffer, T., Tanaka, Y., Tidswell, M., Erle, D.J., Horgan, K.J., Luce, G.E., Lazarovits, A.I., Buck, D., and Shaw, S. (1993). Selective expression of integrin alpha 4 beta 7 on a subset of human CD4+ memory T cells with Hallmarks of gut-trophism. J Immunol 151, 717–729.

Sefik, E., Geva-Zatorsky, N., Oh, S., Konnikova, L., Zemmour, D., McGuire, A.M., Burzyn, D., Ortiz-Lopez, A., Lobera, M., Yang, J., et al. (2015). MUCOSAL IMMUNOLOGY. Individual intestinal symbionts induce a distinct population of RORgamma(+) regulatory T cells. Science 349, 993–997.

Sha, Y., and Markovic-Plese, S. (2011). A role of IL-1R1 signaling in the differentiation of Th17 cells and the development of autoimmune diseases. Self Nonself 2, 35–42.

Shaw, M.H., Kamada, N., Kim, Y.G., and Nunez, G. (2012). Microbiota-induced IL-1beta, but not IL-6, is critical for the development of steady-state TH17 cells in the intestine. J Exp Med 209, 251–258.

Szabo, S.J., Sullivan, B.M., Stemmann, C., Satoskar, A.R., Sleckman, B.P., and Glimcher, L.H. (2002). Distinct effects of T-bet in TH1 lineage commitment and IFN-gamma production in CD4 and CD8 T cells. Science 295, 338–342.

Tanaka, T., and Kishimoto, T. (2012). Targeting interleukin-6: all the way to treat autoimmune and inflammatory diseases. Int J Biol Sci 8, 1227–1236.

Teng, F., Klinger, C.N., Felix, K.M., Bradley, C.P., Wu, E., Tran, N.L., Umesaki, Y., and Wu, H.J. (2016). Gut Microbiota Drive Autoimmune Arthritis by Promoting Differentiation and Migration of Peyer’s Patch T Follicular Helper Cells. Immunity 44, 875–888.

Veldhoen, M., Hocking, R.J., Atkins, C.J., Locksley, R.M., and Stockinger, B. (2006). TGFbeta in the context of an inflammatory cytokine milieu supports de novo differentiation of IL-17-producing T cells. Immunity 24, 179–189.

Wang, C., Yosef, N., Gaublomme, J., Wu, C., Lee, Y., Clish, C.B., Kaminski, J., Xiao, S., Meyer Zu Horste, G., Pawlak, M., et al. (2015). CD5L/AIM Regulates Lipid Biosynthesis and Restrains Th17 Cell Pathogenicity. Cell 163, 1413–1427.

Wilson, N.J., Boniface, K., Chan, J.R., McKenzie, B.S., Blumenschein, W.M., Mattson, J.D., Basham, B., Smith, K., Chen, T., Morel, F., et al. (2007). Development, cytokine profile and function of human interleukin 17-producing helper T cells. Nat Immunol 8, 950–957.

Wu, H.J., Ivanov, II, Darce, J., Hattori, K., Shima, T., Umesaki, Y., Littman, D.R., Benoist, C., and Mathis, D. (2010). Gut-residing segmented filamentous bacteria drive autoimmune arthritis via T helper 17 cells. Immunity 32, 815–827.

Xu, M., Pokrovskii, M., Ding, Y., Yi, R., Au, C., Harrison, O.J., Galan, C., Belkaid, Y., Bonneau, R., and Littman, D.R. (2018). c-MAF-dependent regulatory T cells mediate immunological tolerance to a gut pathobiont. Nature 554, 373–377.

Yang, X.O., Panopoulos, A.D., Nurieva, R., Chang, S.H., Wang, D., Watowich, S.S., and Dong, C. (2007). STAT3 regulates cytokine-mediated generation of inflammatory helper T cells. J Biol Chem 282, 9358–9363.

Yang, Y., Torchinsky, M.B., Gobert, M., Xiong, H., Xu, M., Linehan, J.L., Alonzo, F., Ng, C., Chen, A., Lin, X., et al. (2014). Focused specificity of intestinal TH17 cells towards commensal bacterial antigens. Nature 510, 152–156.

Zhou, L., Ivanov, II, Spolski, R., Min, R., Shenderov, K., Egawa, T., Levy, D.E., Leonard, W.J., and Littman, D.R. (2007). IL-6 programs T(H)-17 cell differentiation by promoting sequential engagement of the IL-21 and IL-23 pathways. Nat Immunol 8, 967–974.

